# Efficient plasmid transfer via natural competence in a synthetic microbial community

**DOI:** 10.1101/2020.10.19.342733

**Authors:** Yu-Yu Cheng, Zhichao Zhou, James M. Papadopoulos, Jason D. Zuke, Tanya G. Falbel, Karthik Anantharaman, Briana M. Burton, Ophelia S. Venturelli

## Abstract

The molecular and ecological factors shaping horizontal gene transfer (HGT) via natural transformation in microbial communities are largely unknown, which is critical for understanding the emergence of antibiotic-resistant pathogens. We investigate key factors shaping HGT in a microbial community by quantifying extracellular DNA release, species growth and HGT efficiency over time. In the community, plasmid release and HGT efficiency are significantly enhanced than the respective monocultures. The donor is a key determinant of HGT efficiency as plasmids induce the SOS response, enter a multimerized state, and are released at high concentrations, yielding efficient HGT. HGT is independent of the donor viability state as live and dead donor cells transfer the plasmid with high efficiency and is only reduced in response to high donor lysis rates. In sum, plasmid HGT via natural competence depends on an interplay of plasmid properties, donor stress responses and lysis rates and inter-species interactions.

## INTRODUCTION

Horizontal gene transfer (HGT) is a major mechanism of genetic variation in microbial communities that enables the acquisition of new functional capabilities^1^. Horizontally acquired sequences can provide a selective advantage by facilitating evolutionary adaptation to changing environmental conditions^2^. Conjugation and natural transformation are prevalent processes that enable HGT in bacterial communities. Conjugation involves cell-to-cell contact between a donor and recipient cell and thus the live donor cell actively participates in the HGT process^3^. By contrast, extracellular DNA (eDNA) can be acquired by a recipient cell that has activated the natural competence pathway in the absence of living donor cells^4^. The capability for natural competence is widespread across Gram-positive and Gram-negative bacteria^5^ and is also a common trait among bacterial pathogens^6^. For example, inter-species or intra-strain gene transfer^7–9^ via natural transformation is implicated in the ability of *Streptococcus pneumonia* (*S. pneumonia*) to adapt and persist on a human host.

eDNA derived from bacteria is prevalent in natural environments^10^. The potential biological functions of eDNA as well as the mechanisms mediating DNA release varies across bacterial species and environmental contexts^11^. eDNA can be released through autolysis, active secretion or cell death and is involved in biofilm formation, DNA repair, nutrient source, and gene transfer^12–15^. Environmental stressors such as bactericidal antibiotics, temperature shifts, chemicals and enzymes can also modify cell viability and thus the rate of eDNA release^16,17^. In addition, interactions between living donors and recipients can influence eDNA release and the frequency of HGT in microbial communities. For instance, naturally competent species can exploit predation to enhance DNA release from donor cells. For example, *Acinetobacter baumannii, Vibrio cholerae* and *S. pneumoniae* can use type-VI secretion systems or bacteriocins to enhance DNA release via lysis of the donor^18–20^. In addition, the presence of a donor strain has been shown to enhance the frequency of gene transfer for intra-strain gene transfer in *B. subtilis, Porphyromonas gingivalis* and *Pseudomonas stutzeri*, as well as inter-species gene transfer between the donor *E. coli* and recipient *Vibrio* species^21–24^. This implies that HGT via natural competence in microbial communities can be mediated by key ecological factors.

At the molecular level, the DNA strand exchange protein and master regulator of the SOS response RecA has been shown to play an important role in facilitating HGT^25^. In the recipient, RecA mediates homologous recombination with DNA with sufficient homology to the recipient genome^26^. In the donor, RecA plays an important role in HGT via transduction and conjugation^27,28^. In response to DNA damage, RecA binds to single-stranded DNA and inactivates the LexA repressor that regulates the SOS response genes (global response to DNA damage)^29^. Depending on the degree of DNA damage, the SOS response activates expression of error-prone DNA polymerases, inhibitors of cell division, and proteins that promote cell death^30^. In addition, RecA triggers the degradation of repressors in phages and integrative and conjugative elements with homology to LexA, which in turn enhances the rates of transduction and conjugation^27,28^. For natural transformation, RecA has been shown to affect the multimerization of plasmids, which have a higher transformation frequency than monomeric plasmids^31,32^. However, the role of RecA in eDNA release and HGT via natural competence in microbial communities are not well understood.

We study dynamics of HGT via natural competence in a synthetic microbial community composed of the donor *E. coli* and recipient *B. subtilis*. We interrogate the contributions of environmental and molecular factors on eDNA release and efficiency of HGT. We show that eDNA release and HGT efficiency via natural competence depends on the initial densities of the donor and recipient, mirroring conjugation^33^. Notably, *recA* harbored by the *E. coli* donor has a major impact on HGT frequencies due to its effects on plasmid multimerization and eDNA release. Using inducible control of cell lysis, we show that high rates of donor lysis inhibit HGT of a plasmid due to competition with genomic DNA (gDNA) and the release of DNases into the environment. By comparison, we demonstrate that live, heat-killed, or antibiotic-inhibited donor cells can efficiently release eDNA which in turn yields high HGT efficiencies in the synthetic microbial community. Bioinformatic analysis identifies *E. coli* plasmid replication origins in wild-type *Bacillus* genomes, suggesting that plasmids may be transferred more broadly to *B. subtilis*. In sum, our results provide key insights into the molecular and environmental factors influencing HGT frequency via natural competence in microbial communities.

## RESULTS

### Inter-species plasmid HGT is efficient in a microbial community

We investigated inter-species HGT mediated by natural competence in a synthetic community composed of the donor *E. coli* and recipient *B. subtilis* (**Fig. 1a**). This synthetic community enabled quantification of time-resolved changes in species abundance, eDNA release and HGT frequency. The *E. coli* donor harbored an integrative plasmid pBB275 with a spectinomycin resistance gene (*specR*) flanked by two ∼500 bp sequences homologous to *B. subtilis* PY79 *ycgO*. HGT occurred when eDNA encompassing the pBB275 integrative plasmid was taken up and integrated via homologous recombination onto the genome of *B. subtilis* with an active natural competence program. In addition, the donor *E. coli* strain (MG1655-rfp) in the co-culture contained a constitutively expressed red fluorescent protein (*rfp*) integrated onto the genome for fluorescent imaging.

**Figure 1.**
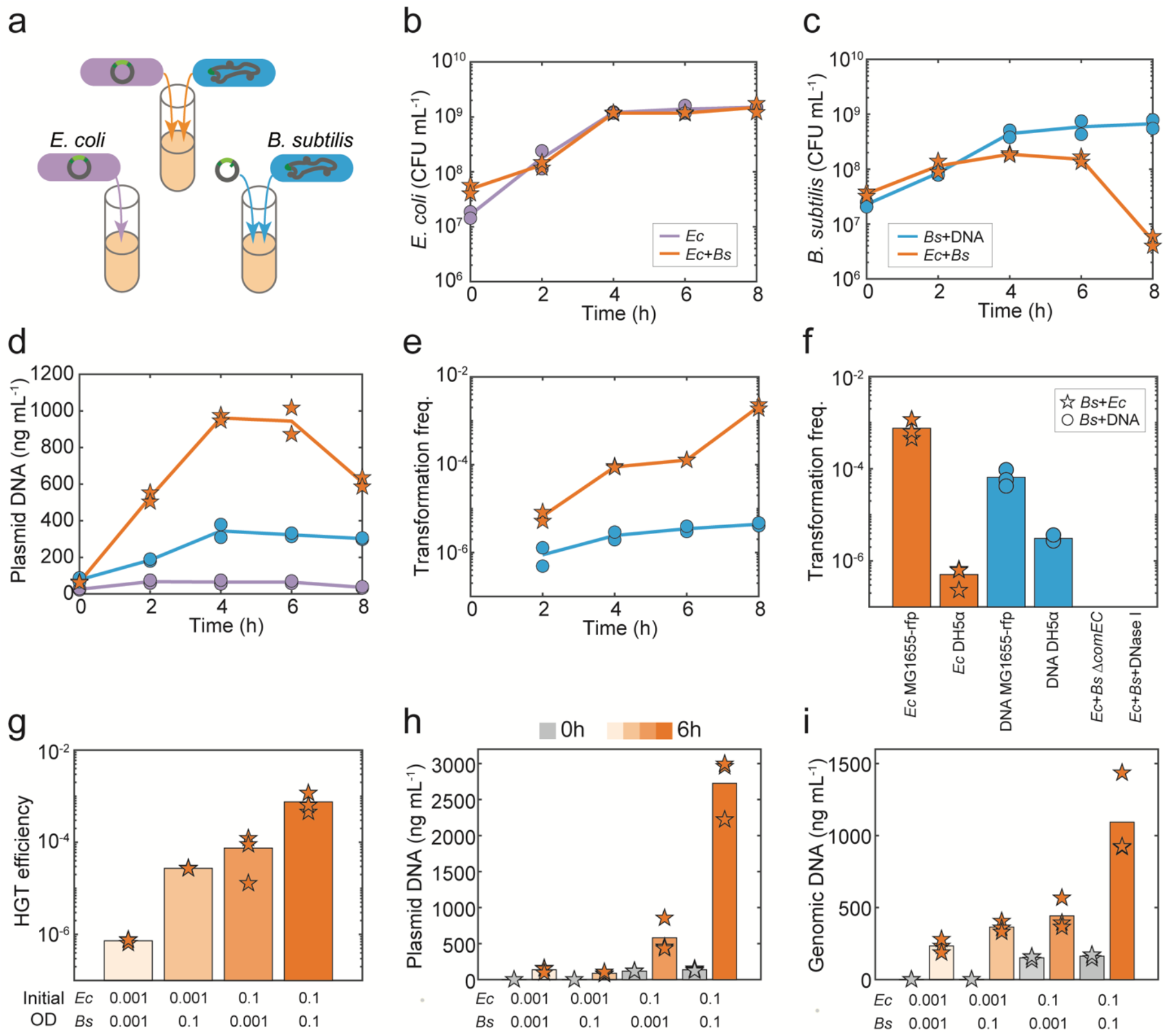
Plasmid horizontal gene transfer is highly efficient in the microbial community. **(a)** Schematic of experimental design to characterize the temporal changes in plasmid horizontal gene transfer (HGT), species abundance and extracellular plasmid release in the xylose-inducible *comK B. subtilis* monoculture supplemented with 100 ng/mL pBB275 plasmid DNA extracted from *E. coli* DH5α (blue) or community composed of *E. coli* MG1655-rfp harboring pBB275 and xylose-inducible *comK B. subtilis* (orange). The pBB275 plasmid harbored a spectinomycin resistance gene flanked by two ∼500 bp *B. subtilis* PY79 *ycgO* sequences homologous to the *B. subtilis* genome. *B. subtilis* was engineered to harbor a xylose-inducible master regulator for competence *comK* for enhancing transformation efficiency. Time-series measurements of **(b)** *E. coli* MG1655-rfp harboring pBB275 abundance, **(c)** xylose-inducible *comK B. subtilis* abundance, **(d)** extracellular pBB275 plasmid concentration, and **(e)** transformation frequency of the pBB275 plasmid in the *B. subtilis* monoculture and community. **(f)** Bar plot of transformation frequency of the pBB275 plasmid at 6 hr in the monoculture or community. *E. coli* MG1655-rfp harboring pBB275 or *E. coli* DH5α harboring pBB275 was used as the donor species in the community. In the monoculture, 1 μg/mL pBB275 plasmid DNA derived from *E. coli* MG1655-rfp or DH5α was introduced into the xylose-inducible *comK B. subtilis* monoculture. The transformation frequencies were below the detection limit of ∼10^−9^ for the community composed of *E. coli* M1655-rfp harboring pBB275 and *B. subtilis* Δ*comEC* and the community composed of *E. coli* MG1655-rfp harboring pBB275 and xylose-inducible *comK B subtilis* in the presence of 1 unit mL^-1^ DNase I. **(g)** HGT efficiency (i.e. transformation frequency in community) of the pBB275 plasmid at 6 hr in communities composed *E. coli* MG1655-rfp pBB275 and xylose-inducible competent *B. subtilis* inoculated at different initial abundances (absorbances at 600 nm or OD). **(h)** Bar plot of measured extracellular pBB275 plasmid concentration in the community composed of *E. coli* MG1655-rfp pBB275 and xylose-inducible *comK B. subtilis*. **(i)** Extracellular genomic *E. coli* MG1655-rfp concentration at 0h or 6h in the community composed of *E. coli* MG1655-rfp pBB275 and xylose-inducible *comK B. subtilis* inoculated at different initial densities. All time-series experiments in (a), (c), (d) and (e) had two biological replicates. All single time measurements in (f), (g), (h) and (i) had three biological replicates. Lines and bars represent the average of the biological replicates.

To understand the dynamics of HGT, we measured the colony forming unit (CFU) of the *E. coli* donor, *B. subtilis* recipient, and *B. subtilis* transformants over time in monoculture and co-culture conditions (**Fig. 1a,b,c**). To investigate the temporal changes in eDNA release, we performed time-series quantitative real-time PCR (qPCR) measurements of the pBB275 plasmid harbored by *E. coli* (MG1655-rfp) in monoculture, co-culture of *E. coli* (MG1655-rfp) and *B. subtilis*, and *B. subtilis* monoculture in the presence of purified pBB275 derived from the widely used cloning strain *E. coli* DH5α (**Fig. 1a,d**). To evaluate the efficiency of HGT in the community, we quantified the transformation frequency as a function of time in the co-culture or *B. subtilis* monoculture containing a high concentration of pBB275 (100 ng/mL) (**Fig. 1a,e**). Transformation frequency was defined as the ratio of the number of transformants plated on selective media to the total number of *B. subtilis*. To enhance transformation efficiency (∼100-fold) in rich media, the *B. subtilis* strain harbored a xylose-inducible master regulator for competence (*comK*)^34^. Based on a titration of plasmid concentration and transformation efficiency of the *B. subtilis* monoculture, we found that 100 ng/mL was close to the saturated regime of the dose response (**Supplementary Fig. 1a**). This indicates that higher plasmid concentrations beyond 100 ng/mL would not substantially enhance transformation frequency.

The growth of *E. coli* was similar in the presence and absence of *B. subtilis* (**Fig. 1b**). However, the growth of *B. subtilis* after 4 hr was reduced in the presence of *E. coli*, indicating a negative inter-species interaction from *E. coli* to *B. subtilis* (**Fig. 1c**). The fluorescently labelled *E. coli* and *B. subtilis* aggregated together, suggesting that the mechanism of *B. subtilis* growth inhibition involved cell-to-cell contact (**Supplementary Fig. 1b**). The growth inhibition was observed for both the wild-type and engineered *B. subtilis* strain (**Fig. 1c and Supplementary Fig. 1c**). In addition, eDNA release from *E. coli* was substantially enhanced in the presence of *B. subtilis* than in the *E. coli* monoculture (**Fig. 1d, orange vs. purple line**). The plasmid concentration moderately increased over time in the *B. subtilis* monoculture, possibly due to autolysis of *B. subtilis*^15^ (**Fig. 1d, blue line**). The transformation frequencies of both the engineered and wild-type *B. subtilis* strains were significantly enhanced in the co-culture than in the respective *B. subtilis* monocultures (**Fig. 1e and Supplementary Fig. 1d**). These data indicate that the presence of *E. coli* substantially enhanced HGT efficiency to *B. subtilis*. This enhancement could be due to higher rates of eDNA release and/or differences in the genetic background of the *E. coli* donor (MG1655 vs. DH5α).

We considered if differences in plasmid concentration and/or multimerization state due to variation in the *E. coli* donor (i.e. purified plasmid derived from DH5α or released plasmid from MG1655-rfp) drove the high transformation frequency observed in co-culture. Indeed, plasmids derived from *E. coli* MG1655-rfp contained multimers (**Supplementary Fig. 1e**). However, plasmids derived from DH5α did not form multimers, consistent with the presence and absence of *recA* multimerization activity in these strains^31^. Consistent with the previously reported higher transformation frequency of multimerized plasmids^32^, *B. subtilis* monoculture supplemented with plasmid DNA (1 μg/mL) from *E. coli* MG1655-rfp had ∼10-fold higher transformation frequency than *B. subtilis* monoculture exposed to the same plasmid concentration derived from *E. coli* DH5α (**Fig. 1f, blue bars**).

We focused on the transformation frequency at 6 hr due to the high abundance of *B. subtilis* and transformation frequency observed in co-culture (**Fig. 1c,e**). Consistent with this result, the transformation frequency of the *B. subtilis* monoculture was consistently ∼10-fold higher in the presence of a broad range of plasmid concentrations derived from *E. coli* MG1655-rfp than DH5α (**Supplementary Fig. 1a**). Notably, the transformation frequency in co-culture with *E. coli* DH5α was ∼3 orders of magnitude lower than in co-culture with *E. coli* MG1655-rfp (**Fig. 1f, orange bars**). This implies that while plasmid multimerization was a key factor that enhanced HGT efficiency, it did not fully explain the magnitude of the enhancement of HGT efficiency observed in co-culture with MG1655 compared to DH5α.

To test the possibility that the plasmid transfer occurred via other mechanisms such as conjugation, we co-cultured *B. subtilis* PY79 Δ*comEC* (transport protein for DNA binding and uptake^35^) with *E. coli* MG1655-rfp. In this community, transformants were not observed at 6 hours (**Fig. 1f**). Consistent with this result, the presence of DNase I abolished plasmid transfer in the co-culture with the engineered *B. subtilis* and *E. coli* MG1655-rfp. In sum, these results indicate plasmid transfer occurred via natural competence in the community.

We varied the initial cell densities of *E. coli* and *B. subtilis* in the co-culture to understand its impact on eDNA release and HGT efficiency. The plasmid eDNA and HGT efficiency increased with the initial cell densities of *E. coli* and *B. subtilis* (**Fig. 1g,h**). Variation in the initial abundance of both species had a similar impact on HGT efficiency, where the highest initial abundance of both species yielded the highest HGT efficiency (**Fig. 1g**). The extracellular gDNA of *E. coli* increased with its initial abundance in the community (**Fig. 1i**), indicating a higher abundance of lysed cells. Lower initial *E. coli* abundance resulted in higher abundance of *B. subtilis* at 6 hr, consistent with *E. coli* ‘s negative impact on the growth of *B. subtilis* (**Supplementary Fig. 1f**). However, *B. subtilis* abundance at 6 hr did not trend with the large variation in HGT efficiency across conditions. In sum, the initial abundance of the donor and recipient was a critical variable shaping HGT efficiency and eDNA release (**Fig. 1g,h**).

To determine if sequences derived from gDNA can transfer from *E. coli* to *B. subtilis* in the community via natural competence, an erythromycin resistance gene flanked by ∼600 bp sequences homologous to the *B. subtilis* PY79 *yvbJ* was introduced into the *E. coli* MG1655 donor strain (**Supplementary Fig. 2a**). Transformants were not observed over 8 hr in the co-culture of the *E. coli* gDNA donor strain and engineered *B. subtilis* (detection limit was 10^−9^) (**Supplementary Fig. 2b**). The *B. subtilis* monoculture exposed to *E. coli* gDNA (100 ng/mL) displayed a low transformation frequency (**Supplementary Fig. 2b**). While eDNA release of the *E. coli* donor strain was moderately higher in co-culture than in the monoculture (**Supplementary Fig. 2c**), this concentration was not high enough to enable efficient HGT. Mirroring the trends in species growth in the plasmid *E. coli* donor co-culture, the growth of *B. subtilis* was inhibited in co-culture with the gDNA *E. coli* donor (**Supplementary Fig. 2d,e**). In sum, our results indicate that plasmid HGT was substantially more efficient than HGT of sequences derived from gDNA via natural competence in the community.

### Investigating the role of *recA* in plasmid HGT via natural competence

We investigated the factors influencing the ∼10^3^-fold increase in HGT efficiency in the community with *E. coli* MG1655-rfp compared to DH5α (**Fig. 1f, orange bars**). *E. coli* MG1655 and DH5α contain many genetic differences including the presence and absence of the master regulator of the SOS response *recA*, respectively^36,37^. We introduced individual *E. coli* strains that varied in the presence of *recA* (*E. coli* MG1655-rfp or MG1655), absence of *recA* (MG1655 Δ*recA* or DH5α) or constitutively expressed *recA* (DH5α + *recA*) into the community (**Fig. 2a**). Our results showed that donor strains harboring *recA* displayed substantially higher HGT efficiency in the community than *recA*-donors (**Fig. 2a**). Notably, the presence of constitutively expressed *recA* yielded a ∼100-fold increase in HGT efficiency, indicating that *recA* was a major determinant of HGT efficiency. Consistent with this result, plasmid eDNA was higher in communities with *E. coli recA*+ donors than *recA*-donors (**Fig. 2b**). In co-culture, all *E. coli* strains achieved similar cell density at 6 hr (**Supplementary Fig. 3a**). These strains displayed similar doubling times in monoculture, with MG1655-rfp and DH5α + *recA* displaying the fastest and slowest growth rates, respectively (**Supplementary Fig. 3b,c**). In addition, *E. coli recA*+ donors displayed higher extracellular gDNA than *recA*-donors in the community (**Fig. 2c**), suggesting that RecA can facilitate plasmid transfer by increasing the rate of cell death and eDNA release. Constitutively expressed *rfp* in *E. coli* MG1655 enhanced eDNA release, suggesting that the metabolic burden imposed by heterologous gene expression enhanced the rate of cell death (**Fig. 2b,c**).

**Figure 2.**
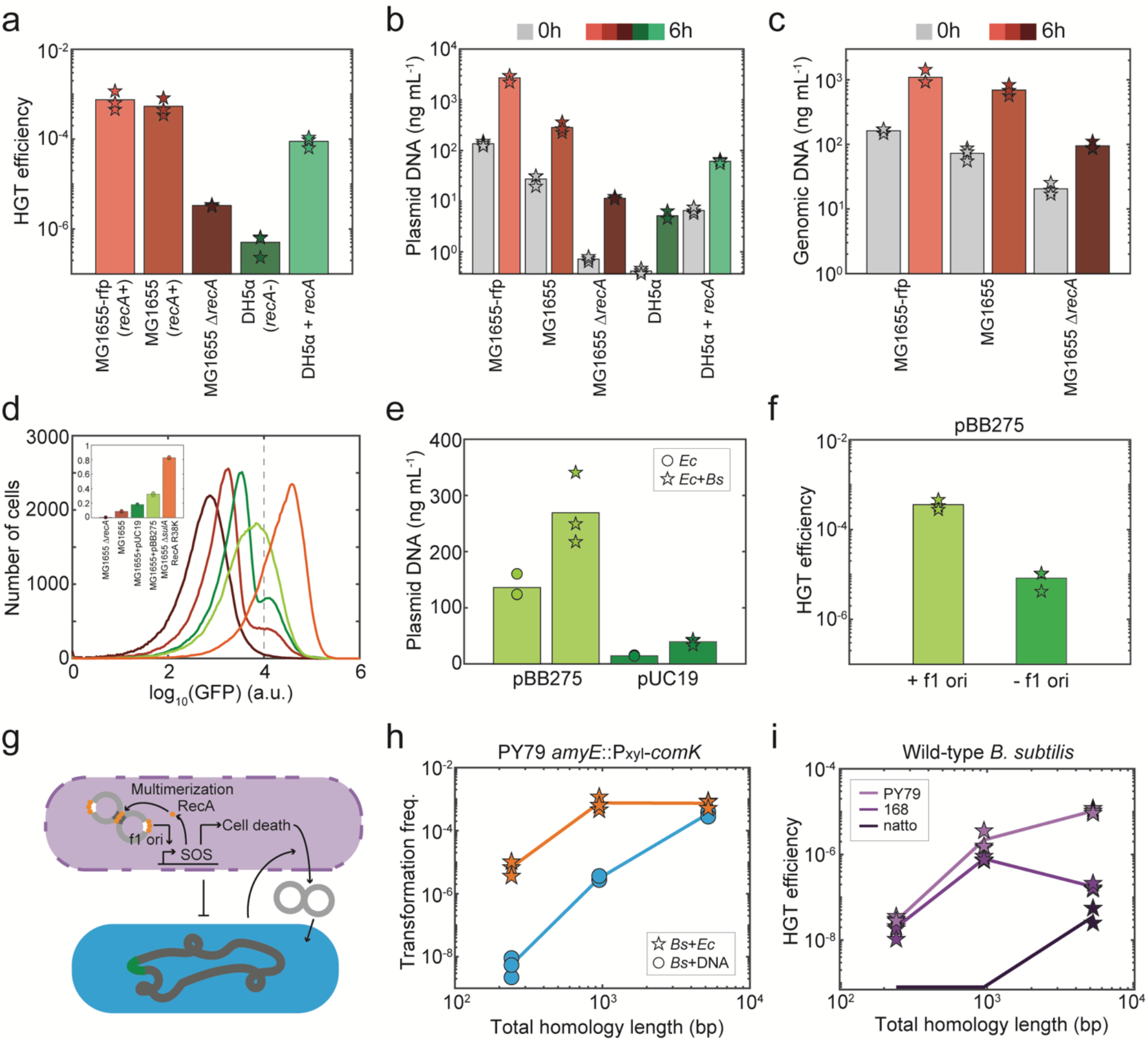
The presence of *recA* in the *E. coli* donor enhances HGT efficiency in the community. **(a)** Bar plot of HGT efficiencies of the pBB275 plasmid in the community with an *E. coli recA*+ or *recA*-donor and xylose-inducible *comK B. subtilis* at 6 hr. **(b)** Extracellular pBB275 plasmid concentration or **(c)** extracellular genomic DNA concentration in communities composed of a *recA*+ or *recA*-*E. coli* plasmid donor and xylose-inducible *comK B. subtilis* at 0 or 6 hr. **(d)** Histogram of GFP expression driven by a SOS response promoter in *E. coli* MG1655 Δ*recA, E. coli* MG1655, *E. coli* MG1655 harboring pUC19, *E. coli* MG1655 harboring pBB275 or *E. coli* MG1655 Δ*sulA* RecA E38K in monoculture measured by flow cytometry. Inset: bar plot of the fraction of *E. coli* expressing high GFP (>10000 a.u.) for each strain. **(e)** Extracellular plasmid concentration in the *E. coli* MG1655 pBB275 or pUC19 monoculture or community with xylose-inducible *comK B. subtilis*. **(f)** HGT efficiencies in the community composed of *E. coli* MG1655 pBB275 with or without a f1 ori and xylose-inducible *comK B. subtilis*. **(g)** Schematic of proposed model for the efficient plasmid transfer via natural competence in a microbial community composed of a *recA*+ *E. coli* donor and recipient *B. subtilis*. The plasmid leads to the activation of the SOS response, which in turn enhances eDNA release. RecA facilitates plasmid multimerization. *B. subtilis* enhances *E. coli* eDNA release and *E. coli* inhibits the growth of *B. subtilis*. **(h)** Relationship between homology length and the transformation frequency of the pBB275 plasmid in the community composed of *E. coli* MG1655-rfp pBB275 and xylose-inducible *comK B. subtilis* or *B. subtilis* monoculture at 6 hr. In the *B. subtilis* monoculture, 1 μg/mL pBB275 plasmid DNA derived from *E. coli* DH5α was introduced. The pBB275 plasmid in *E. coli* MG1655-rfp had three different total homology lengths - ∼200 bp, ∼1000 bp, and ∼5000 bp. **(i)** Relationship between homology length and HGT efficiency of the pBB275 plasmid in a community composed of the DAP-auxotrophic *E. coli* BW29427 pBB275 and wild-type *B. subtilis* PY79, 168 or natto IFO3335 at 6 hr. Transformants were selected with 100 μg/mL spectinomycin in the absence of DAP. HGT efficiency of pBB275 plasmid with total ∼200 bp and ∼1000 bp homology for *B. subtilis* natto was below the detection limit ∼10^−9^. All experiments had three biological replicates. Bars and lines are the average of the biological replicates.

RecA can control the rate of cell death via activation of the SOS response^38^. Therefore, we tested if the integrative plasmid pBB275 induced the SOS response, which in turn could enhance the rate of cell death and eDNA release. To quantify the activity of the SOS response, we constructed the SOS response reporter P_sulA_-*gfp* on a second plasmid^39^. We introduced the SOS response reporter plasmid into *E. coli* MG1655 Δ*recA*, MG1655, and MG1655 Δ*sulA* RecA E38K, which has a constitutively active SOS response^40^. Plasmid pBB275 consists of heterologous sequences including the f1 replication origin (f1 ori), a remnant of the construction of the original plasmid (**Supplementary Fig. 4a**). Heterologous sequences as well as the f1 ori have been shown to activate the SOS response in *E. coli*^41–43^.

We used flow cytometry to quantify the distribution of the SOS GFP reporter across the population. Our results showed that 1% of *E. coli* MG1655 Δ*recA*, 9% of *E. coli* MG1655, and 83% of wild-type *E. coli* MG1655Δ*sulA* RecA E38K displayed high GFP fluorescence, indicating that this GFP ON sub-population had activated the SOS response (**Fig. 2d**). To evaluate the effect of the pBB275 plasmid on the activity of the SOS response, we introduced a plasmid that lacks the f1 and contains the *lacZα* gene (pUC19) into *E coli* MG1655 (**Supplementary Fig. 4b**). *E. coli* MG1655 harboring pUC19 or pBB275 displayed larger GFP ON sub-populations (18% and 33% of the populations, respectively) than MG1655 lacking these plasmids (9% of the population). The GFP ON sub-population of *E. coli* MG1655 harboring pBB275 displayed elongated cell morphologies based on fluorescent microscopy imaging compared to the GFP ON sub-population of the same strain harboring pUC19 (**Supplementary Fig. 5a-d**).

In monoculture and in the community, plasmid eDNA release was substantially higher for the *E. coli* MG1655 harboring pBB275 than *E. coli* MG1655 harboring pUC19 (**Fig. 2e**). This is consistent with the observed larger fraction of the SOS reporter GFP ON cells in *E. coli* with pBB275 than pUC19 (**Fig. 2d**). In addition, the plasmid eDNA concentration was higher in communities containing both individual *E. coli* strains than in their respective monocultures, consistent with the trends in *E. coli* MG155-rfp (**Fig. 1d and 2e**). In the absence of f1 ori and sequences homologous to *B. subtilis*, the conformations of pUC19 purified from *recA*+ *E. coli* or *recA*-*E. coli* were similar (**Supplementary Fig. 5e**). To determine if other sequence differences between pUC19 and pBB275 contributed to the observed differences in HGT efficiency and plasmid eDNA release, we constructed a derivative of pBB275 that lacks the f1 ori (**Supplementary Fig. 4c**). Our results demonstrated that the HGT efficiency of pBB275 containing the f1 ori was significantly higher than *E. coli* harboring pBB275 lacking the f1 ori in the community, indicating that the f1 ori was a critical determinant of HGT efficiency (**Fig. 2f**). These results suggest that the f1 ori may activate the SOS response and enhance plasmid multimerization and plasmid eDNA release via RecA, which in turn enhances HGT efficiency (**Fig. 2g**). In the presence of a high concentration of multimeric pBB275 (1 μg/mL purified from MG1655-rfp), HGT efficiency was higher in the co-culture than monoculture (**Fig. 1f**). This difference suggests that the inter-species interactions inhibiting *B. subtilis* growth and enhancing *E. coli* eDNA release may lead to higher HGT efficiency (**Fig. 1c, Supplementary Fig. 1c,f and 2e**).

To understand how other plasmid features impacted HGT efficiency, we studied the relationship between the length of homology (∼200bp, ∼1000bp, or ∼5000bp) to *B. subtills* and HGT efficiency, eDNA release and plasmid multimerization using pBB275 containing the f1 ori. Plasmids with different homology lengths all formed multimers in *E. coli* MG1655-rfp (*recA*+) (**Supplementary Fig. 5f**).The HGT efficiency was substantially higher in the community than in monoculture exposed to a high concentration of these plasmids (1 μg/mL purified from DH5α) (**Fig. 2h**). The magnitude of this enhancement in HGT efficiency was maximized (∼1000-fold) for the plasmid with the shortest homology length (**Fig. 2h**). The plasmid eDNA concentration displayed a non-monotonic relationship with homology length and was maximized for ∼1000 bp homology (**Supplementary Fig. 5g**). This implies that the length of the homology influenced the rate of cell death and thus eDNA release.

To demonstrate that plasmids with homology to *B. subtilis* can transfer easily from *recA*+ *E. coli* to *B. subtilis* in the community, we co-cultured three individual wild-type *B. subtilis* strains including PY79, 168, and natto IFO3335, with *E. coli* BW29427, a *recA*+ and diaminopimelic acid (DAP) auxotrophic strain^44–46^. By selecting for spectinomycin resistant cells in the absence of DAP, we found that HGT can occur in the presence of all individual *B. subtilis* strains in their corresponding pairwise communities (**Fig. 2i**). Therefore, our results suggest that the plasmids can transfer efficiently from *E. coli* donor strains to different *B. subtilis* strains via natural competence.

### High donor cell lysis rates and DNA competition reduces HGT efficiency

Bacteria are continuously confronted with biotic and abiotic environmental stimuli that impact cellular growth rates and viability^47,48^. We investigated how donor growth perturbations impacted HGT efficiency in synthetic communities. To precisely control the lysis rate of the donor, we introduced an Isopropyl β-D-1-thiogalactopyranoside (IPTG)-inducible phage φX174 lysis gene *E* into the *E. coli* MG1655-rfp donor strain on a second plasmid in addition to the plasmid pBB275 (**Fig. 3a**). The *E. coli* abundance decreased with IPTG concentration at 6 hr, consistent with the inhibition of growth by the lysis gene (**Fig. 3b**). In addition, *B. subtilis* grew to a moderately higher abundance as a function of IPTG concentration, consistent with a weakened negative interaction from *E. coli* to *B. subtilis* (**Fig. 3b**). The concentration of eDNA for plasmid and gDNA were substantially higher in the presence of IPTG, consistent with the reduced abundance of *E. coli* (**Fig. 3c-f**). Notably, despite the higher eDNA concentrations in the presence of IPTG, the HGT efficiency and abundance of *B. subtilis* transformants decreased over a range of IPTG concentrations and then increased at the highest IPTG concentration (**Fig. 3g,h**).

**Figure 3.**
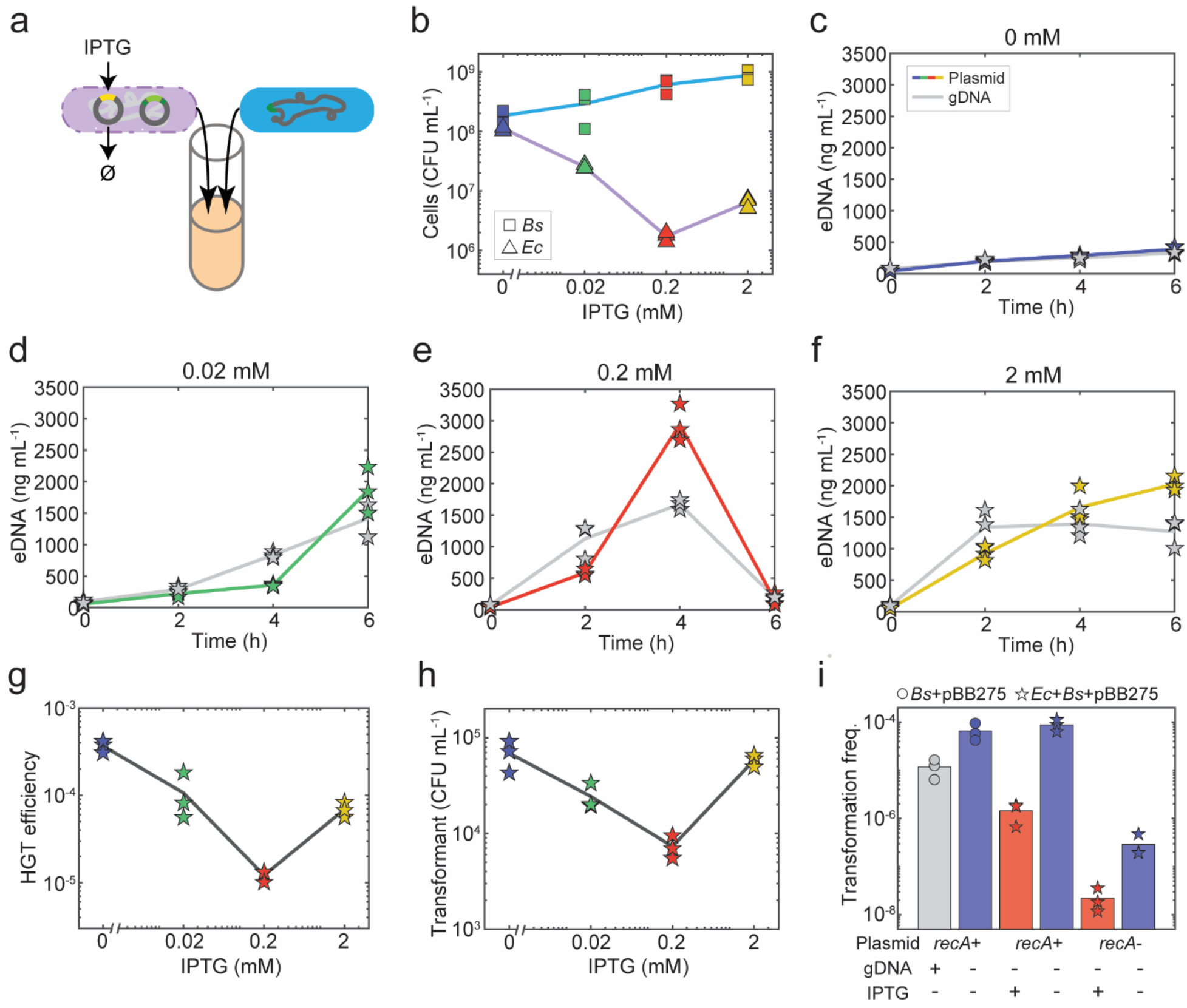
High *E. coli* donor cell lysis rates inhibits HGT efficiency in the community. **(a)** Schematic of the community experiment composed of xylose-inducible *comK B. subtilis* and *E. coli* MG1655-rfp harboring pBB275 and an IPTG-inducible lysis gene. **(b)** *E. coli* and *B. subtilis* abundances in the community in the presence of 0, 0.02, 0.2 or 2 mM IPTG at 6 hr. Time-series measurements of extracellular plasmid or *E. coli* gDNA concentrations in the community in the presence of **(c)** 0 mM, **(d)** 0.02 mM, **(e)** 0.2 mM or **(f)** 2 mM IPTG. **(g)** HGT efficiency of the pBB275 plasmid in the community in the presence of 0, 0.02, 0.2 or 2 mM IPTG at 6 hr. **(h)** Abundance of transformants at 6 hr in the community in the presence of 0, 0.02, 0.2 or 2 mM IPTG. **(i)** Bar plot of transformation frequencies of xylose-inducible *comK B. subtilis* in monoculture or community with *E. coli* harboring an IPTG-inducible lysis gene. In the *B. subtilis* monoculture, 1 μg/mL pBB275 plasmid DNA derived from *E. coli* MG1655-rfp (*recA*+) or DH5α (*recA*-) was introduced. The presence of 1 μg/mL genomic DNA derived from *E. coli* MG1655-rfp or 0.2 mM IPTG in the community reduced the transformation frequency of the pBB275 plasmid DNA at 6h. Each experiment had three biological replicates. Lines and bars are the average of the biological replicates.

To test whether the released *E. coli* gDNA may compete with the plasmid and thus reduce transformation efficiency, we introduced a high concentration of *E. coli* MG1655-rfp gDNA (1 μg/mL) into the *B. subtilis* monoculture that also contained a high concentration of pBB275 plasmid DNA (1 μg/mL). The presence of *E. coli* gDNA inhibited transformation frequency in the *B. subtilis* monoculture, suggesting that gDNA competes with the plasmid for DNA uptake and/or limiting homologous recombination machinery (**Fig. 3i, first two bars**). To test whether cell lysis was an additional factor that inhibits HGT of plasmid, we co-cultured *B. subtilis* with an *E. coli* MG1655-rfp strain harboring a single plasmid with the IPTG-inducible lysis gene. We introduced a high concentration of pBB275 (1 μg/mL) purified from DH5α (monomeric) or MG1655-rfp (multimeric) into separate cultures. The transformation frequency was substantially reduced in the presence of IPTG (0.2 mM), suggesting that donor cell lysis reduced the transformation frequency of both monomeric and multimeric plasmids (**Fig. 3i, last four bars**). The pBB275 plasmid added externally was degraded in the supernatants of the co-cultures containing *B. subtilis* and the IPTG-inducible lysis *E. coli* MG1655-rfp at 6 hr (**Supplementary Fig. 6**). This implies that DNases released from lysed *E. coli* could also inhibit plasmid HGT. In sum, these data indicate that competition with other DNA molecules and high donor cell lysis rates substantially reduced HGT efficiency in the community. The non-monotonic trend in the abundance of *B. subtilis* transformants could be attributed to an interplay of inhibition of plasmid HGT efficiency due to DNA competition and release of DNases and enhanced *B. subtilis* abundance as a consequence of high *E. coli* lysis rates (**Fig. 3b,h**).

### Live or dead donor cells display efficient plasmid transfer

Since natural microbial communities consist of both live and dead cells^47,48^, we tested the effect of donor cell viability on HGT efficiency in the community. We observed similar trends in plasmid and gDNA release for heat-killed *E. coli* (60°C for 30 minutes^49^) and *E. coli* that was not heat-treated individually co-cultured with *B. subtills* (**Supplementary Fig. 7a,b)**. This indicates that heat treatment did not induce substantial *E. coli* lysis. The log transform of the HGT efficiency in the co-culture of heat-killed *E. coli* and *B. subtilis* was linearly related to the log transform of the initial density of heat-treated *E. coli* (**Fig. 4a,b**). In addition, the extracellular plasmid concentration increased with the initial density of heat-killed *E. coli* (**Fig. 4c**). This suggests that plasmids can be efficiently released by intact and dead *E. coli* cells.

**Figure 4.**
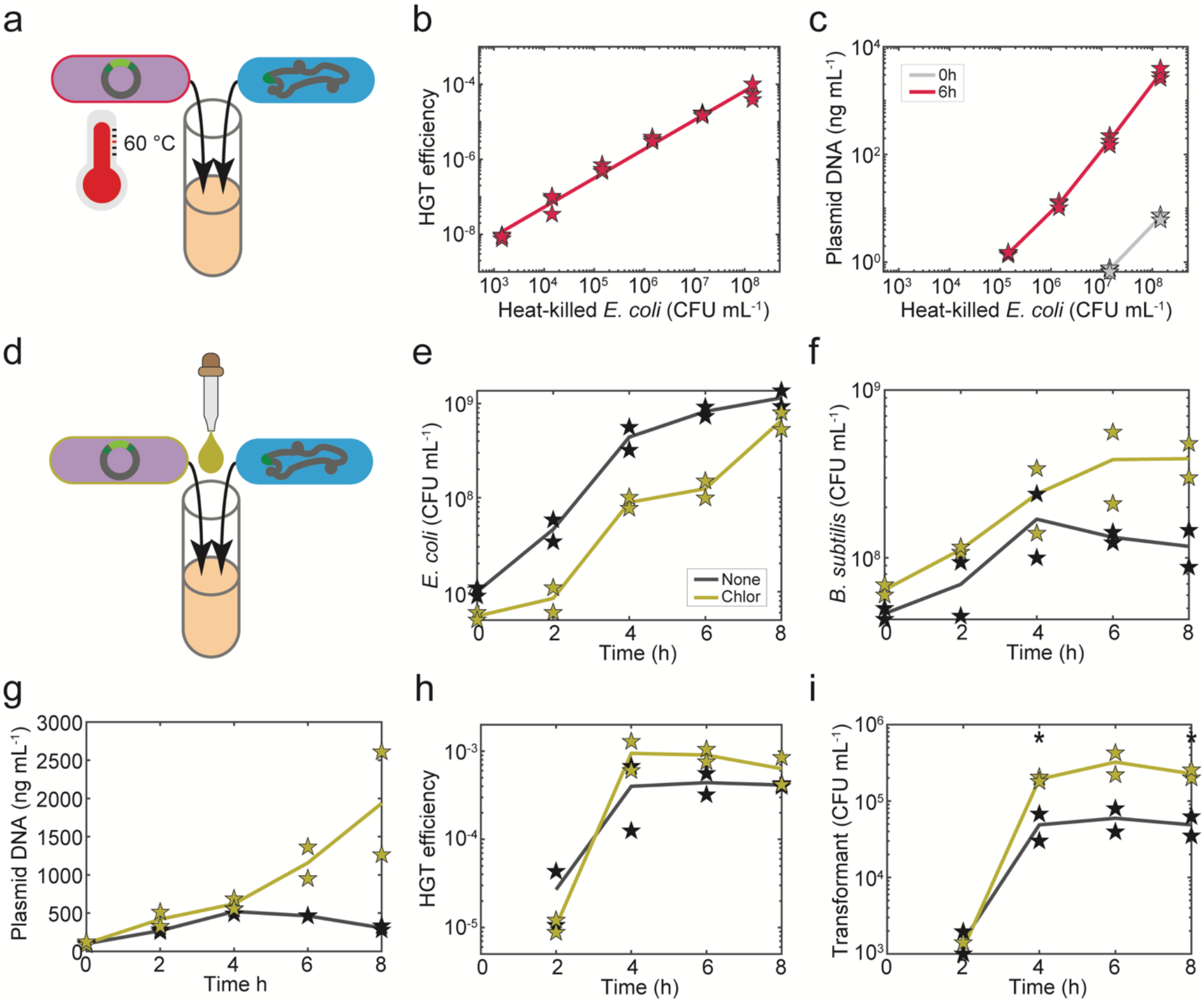
High HGT efficiency in the microbial community is independent of *E. coli* donor viability state. **(a)** Schematic of the experimental design to determine the impact of donor species viability on HGT efficiency and plasmid DNA release. The microbial community was composed of xylose-inducible *comK B. subtilis* and live or heat-killed *E. coli* MG1655-rfp harboring pBB275. **(b)** Scatter plot of the initial abundance of heat-killed *E. coli* MG1655-rfp harboring pBB275 and HGT efficiency in the microbial community at 6 hr. Line represents data fit to the linear equation *y* = 0.774\**x* - 10.37 with the coefficient of determination *R*^2^ = 0.9839 where *x* and *y* are log10 transformed abundance of heat-killed *E. coli* density and the HGT efficiency of pBB275 plasmid, respectively. **(c)** Scatter plot of the initial abundance of heat-killed *E. coli* MG1655-rfp pBB275 and extracellular pBB275 plasmid concentration at 0h or 6 hr in the community. **(d)** Schematic of the experimental design to determine the impact of chloramphenicol (5 μg/mL) on HGT efficiency and plasmid DNA release in the microbial community composed of xylose-inducible *comK B. subtilis* and *E. coli* MG1655-rfp pBB275. *B. subtilis* harbored a chloramphenicol resistance gene. Time-series measurements of **(e)** the abundance of *E. coli* MG1655-rfp harboring pBB275, **(f)** the abundance of xylose-inducible *comK B. subtilis*, **(g)** extracellular pBB275 plasmid concentration, **(h)** HGT efficiency of the pBB275 plasmid, and **(i)** abundance of transformants in the community in the presence or absence of chloramphenicol. Stars (*) denote unpaired *t*-test with *p*-value of 0.0237 and 0.0283 for 4 hr and 8 hr, respectively in the presence and absence of chloramphenicol. Time-series measurements had two biological replicates. Single time measurements had three biological replicates. Lines denote the average of the biological replicates.

Since antibiotics inhibit the growth, viability and lysis rates of bacteria, we introduced a sub-lethal concentration of chloramphenicol (5 μg/mL) into the community that inhibits *E. coli* growth by blocking protein synthesis (**Fig. 4d**). *B. subtilis* harbored a chloramphenicol resistance gene and thus its growth was not significantly impacted by the presence of the antibiotic. In the presence of chloramphenicol, the growth of *E. coli* was reduced, whereas *B. subtilis* abundance was enhanced, consistent with a diminished negative inter-species interaction impacting *B. subtilis* (**Fig. 4e,f**). We found that plasmid release was enhanced in the presence of the antibiotic^50^ (**Fig. 4g**). In addition, extracellular gDNA concentration was higher at 6 hr in the presence than absence of chloramphenicol, suggesting that the antibiotic increased the rate of cell death (**Supplementary Fig. 7c**). HGT efficiency was not significantly higher in the presence of chloramphenicol, consistent with the similar eDNA release in the presence and absence of the antibiotic over the first four hours (**Fig. 4g,h**). However, the abundance of transformants was higher in the presence of chloramphenicol, reflecting the higher abundance of *B. subtilis* in the presence than absence of the antibiotic (**Fig. 4f,i**). The higher *B. subtilis* abundance in this condition is consistent with a reduced abundance in *E. coli* in the presence of chloramphenicol, which in turn diminished the negative inter-species interaction impacting *B. subtilis*. In sum, donor cells, either live or dead, can lead to efficient HGT in microbial communities.

### Plasmid replication origins in wild-type *Bacillus* and non-*Bacillus* genomes

The efficient plasmid transfer in the community suggests that non-conjugative plasmids can efficiently transfer to naturally competent bacteria. To determine if there was a signature of HGT in bacterial genomes, we searched for the ColE1 nucleotide sequence in NCBI Reference Sequence Database (RefSeq) by nucleotide BLAST and found that 31 *Bacillus* strains (77 *Bacillus* hits) harbored ColE1 with high coverage and similarity (>80% coverage and >90% identity) (**Supplementary Fig. 8a and Supplementary Table 1**). Further analyses of the sequence length greater than a threshold length (10^6^ bp) revealed the presence of ColE1 in 6 *Bacillus* genomes (**Fig. 5a**). Notably, other plasmid origins of replication including p15A and CloDF13 did not return as many hits with high percent identity and percent coverage and no hits were observed for pSC101 (**Supplementary Fig. 8b,c and Supplementary Table 1**).

**Figure 5.**
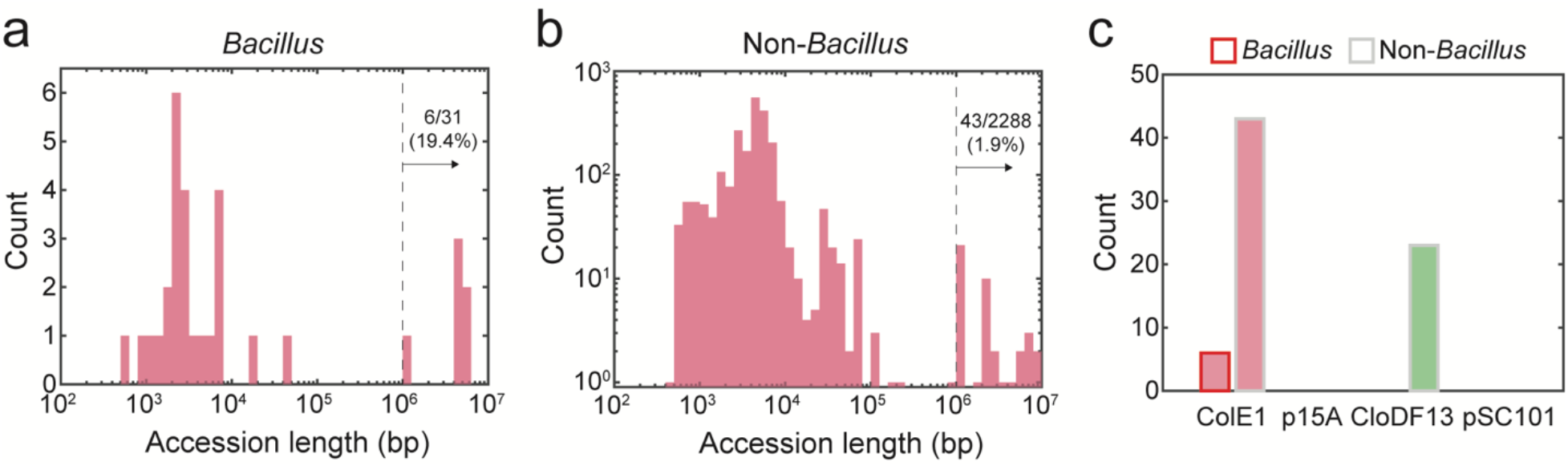
Genomic sequence analysis identifies *E. coli* plasmid replication origins in bacterial genomes. **(a)** Histogram of the sequence lengths of 31 *Bacillus* containing a ColE1 replication origin. **(b)** Histogram of the sequence lengths of 2288 strains excluding members of *Bacillus* with ColE1 replication origin. **(c**) Histogram of the number of *E. coli* plasmid replication origins ColE1, p15A, CloDF13 or pSC101 identified in *Bacillus* or non-*Bacillus* DNA sequences with a sequence length larger than 10^6^ bp. ColE1 was found in 6 *Bacillus* genomes and 43 non-*Bacillus* genomes. CloDF13 was found in 23 non-*Bacillus* genomes. p15A and pSC101 were not detected in this dataset.

A similar analysis for non-*Bacillus* strains revealed that ColE1 and CloDF13 are present in the genomes of 43 and 23 diverse bacteria, respectively (**Fig. 5b,c, Supplementary Fig. 8d-j and Supplementary Table 1**). ColE1 was found in diverse wild-type environmental isolates and laboratory strains, including *Neisseria meningitidis*, which is known to be naturally competent^51^ (**Supplementary Fig. 9a**). By contrast, CloDF13 was found mostly in *Klebsiella pneumoniae* (*K. pneumoniae*) (**Supplementary Table 1**), which may have occurred via conjugation or transduction^52^. A more detailed sequence analysis revealed that ColE1 was associated with antibiotic resistance genes and heterologous proteins in the genomes of *Bacillus* strains and not associated with any known conjugation proteins (**Supplementary Fig. 9b**). In sum, the presence of plasmid replication origins in the genomes of known naturally competent bacteria suggests the possibility that inter-species plasmid transfer has occurred via natural competence in other species and different environments.

## DISCUSSION

Here we used two-member synthetic community composed of *E. coli* and *B. subtilis* to characterize the impact of molecular and environmental factors shaping HGT via natural competence. The transformation frequency of plasmid can be highly efficient in the community and depends on the initial density of both the donor and recipient, mirroring the density-dependence of the rate of conjugation^53^. The phage replication origin f1 was shown to enhance the efficiency of HGT in the community. Future work is needed to understand the role of the f1 ori on the SOS response, and whether the mechanism is direct by potentially forming single-stranded DNA and binding with RecA or indirect^30^. In addition, future work will study the contribution of heterologous genes on the plasmid to plasmid multimerization and HGT efficiencies^41–43^. For example, a recent study indicates that the replication-transcription conflict can lead to plasmid multimerization^54^.

In the community, eDNA release and plasmid HGT efficiency were consistently enhanced compared to the monoculture conditions, indicating that the inter-species interactions influenced HGT efficiency. Previous studies have shown that HGT can be facilitated when the donor and recipient are in close physical contact^21,55^. Here we observed a cell-to-cell interaction between *B. subtilis* and *E. coli* and inhibition of *B. subtilis* growth in the presence of *E. coli*^55,56^. Future studies could explore the effects of cell-to-cell contact on *E. coli* eDNA release, acquisition of eDNA by *B. subtilis* and the growth rate of *B. subtilis*.

Previous studies have investigated the regulation of natural competence in *B. subtilis* to transformation efficiency in monoculture^57^. Here we focused on the molecular and ecological contributions of the donor as opposed to the recipient in shaping HGT. The donor and recipient populations display single-cell heterogeneity in the activation of the SOS response and natural competence program, respectively. Therefore, time-lapse fluorescence microscopy could provide deeper understanding of the role of the donor and recipient in mediating the dynamics of HGT at the single-cell level^18^. In addition, future studies could investigate the spatiotemporal variation in eDNA release using time-lapse microscopy. Our bioinformatic analysis suggests that plasmids could transfer between genetically diverse species via natural competence beyond our synthetic community. The broad transfer of plasmids in microbial communities has implications for the transfer of antibiotic resistance genes^58^. Future work could test more broadly if naturally competent bacteria can take up plasmids from other constituent community members and how plasmid multimerization influences HGT efficiencies^32^.

We show that plasmid transfer from live or intact dead donor cells was highly efficient. However, the efficiency of transfer was reduced in the presence of substantial donor cell lysis. These results suggest that there is an optimal donor lysis rate for maximizing HGT efficiency in microbial communities that can be tuned by the induction of key stress response networks in the donor species. Therefore, we should consider the similar impact of antibiotics on donor cell lysis in addition to the impact of antibiotics on the induction of the natural competence program^27,28^. In sum, a deeper understanding of the molecular networks in the donor strain and environmental and ecological factors influencing HGT in microbial communities could inform microbiome interventions that aim to either reduce HGT frequencies mediating transfer of antibiotic resistance or virulence genes or enhance HGT of key target genes for in situ microbiome engineering^59^.

## MATERIALS AND METHODS

### Plasmids and bacterial strains

To construct the integrative plasmid pBB275, two ∼500 bp *B. subtilis* PY79 *ycgO* sequences were cloned to the upstream and downstream of a spectinomycin resistance gene (*specR*). Two other pBB275 plasmids with ∼100 bp and ∼2500 bp *ycgO* homology were also cloned. pUC19 plasmid was purchased from New England Biolabs. The integration cassette (two *ycgO* homology arms and *specR*) on the pBB275 plasmid was cloned onto pUC19 to replace the *lacZα* gene as the plasmid pBB275 (no f1 ori). To introduce a constitutively expressed RecA into *E. coli* DH5α, the *recA* gene was PCR amplified from *E. coli* MG1655 gDNA and cloned into a p15A plasmid as plasmid pBbA6k_J23100_recA, where RecA is expressed from P_J23100_ promoter. The SOS response reporter plasmid pBbA6c_PsulA_sfGFP was constructed by introducing the 69 bp P_sulA_ promoter to the upstream of green fluorescent protein gene *sfgfp*. The lysis plasmid pYC01 was modified from plasmid pCSaE500, a gift from Lingchong You (Addgene Plasmid #53182). The P_ampC_ promoter of the phage φX174 lysis gene *E* on the pCSaE500 plasmid was replaced with the P_A1lacO-1_ promoter^60^. Plasmids used in this study are listed in **Supplementary Table 2**. To construct the *E. coli* MG1655-rfp, a constitutively expressed P_J23100_-*rfp* was knocked into *E. coli* MG1655 *caiE* locus using the CRISPR gene editing technique^61^. To construct the *E. coli* gDNA donor, an erythromycin resistance gene (*ermR*) flanked by two ∼600 bp *B. subtilis* PY79 *yvbJ* sequence was knocked into the *E. coli* MG1655 *caiE* locus using the same CRISPR method. *E. coli* MG1655 Δ*recA* and *E. coli* MG1655 Δ*sulA* RecA E38K are gifts from Michael Cox. To select transformants of wild-type *B. subtilis*, the DAP-auxotrophic *E. coli* BW29427 (*E. coli* Genetic Stock Center, CGSC) was used as the plasmid donor. Growth of *E. coli* BW29427 required 25 μM 2,6-Diaminopimelic acid (DAP, MilliporeSigma) supplemented in LB. *B. subtilis* PY79 was introduced a xylose-inducible *comK* to increase its transformation efficiency in LB medium^34^. To image the *B. subtilis* under microscope, a constitutively expressed P_hyperspank_-*gfp*(Sp) was integrated into *ycgO* locus^62^. Additional antibiotic resistance genes *ermR* and *kanR* were introduced into *B. subtilis* PY79 for *E. coli* plasmid and gDNA transformation, respectively. *B. subtilis* PY79 was genetically modified using MC medium^63^. MC medium is composed of 10.7 g/L potassium phosphate dibasic (Chem-Impex International), 5.2 g/L potassium phosphate monobasic (MilliporeSigma), 20 g/L glucose (MilliporeSigma), 0.88 g/L sodium citrate dihydrate (MilliporeSigma), 0.022 g/L ferric ammonium citrate (MilliporeSigma), 1 g/L Oxoid casein hydrolysate (Thermo Fisher Scientific), 2.2 g/L potassium L-glutamate (MilliporeSigma), and 20 mM magnesium sulfate (MilliporeSigma). A double crossover of the plasmid into *B. subtilis* PY79 was confirmed by the replacement a different antibiotic resistance gene at the integration locus. gDNA of modified *B. subtilis* was then extracted and transformed into another modified *B. subtilis* to introduce multiple modifications into the genome. Bacterial strains used in this study can be found in **Supplementary Table 3**.

### HGT experiments

To start a monoculture or co-culture of bacteria for transformation, bacteria were first inoculated from the -80 °C glycerol stock into 4 mL Lennox LB media (MilliporeSigma) with selective antibiotics and cultured at 37 °C with shaking (250 rpm) for 12 hours. OD of the overnight culture was measured by NanoDrop One (Thermo Fisher Scientific) and cell culture were diluted to OD0.1 in 5 mL LB in 14 mL Falcon™ Round-Bottom Tube (Thermo Fisher Scientific). LB was supplemented with 50 mM xylose (Thermo Fisher Scientific) to induce *B. subtilis* competence. *E. coli* plasmid donor or gDNA donor carried an antibiotic resistant gene flanked by two *B. subtilis* homology arms, either on the plasmid or in genome. Purified DNA was added to *B. subtilis* monoculture for transformation as a comparison. Cells were well-mixed and then cultured at 37 °C with shaking (250 rpm). Plasmids were extracted using Plasmid Miniprep Kit (Qiagen). gDNA was extracted using DNeasy Blood & Tissue Kit (Qiagen). Cell culture was collected at specific times for qPCR and CFU counting. To quantify eDNA, cells were spun down at 5000 g for 5 min. Then the supernatant was collected and filtered by 0.2 μm Whatman Puradisc Polyethersulfone Syringe Filter (GE Healthcare), stored at -20 °C, and later quantified by qPCR. For CFU counting of donor and recipient, cell culture were serially diluted in phosphate-buffered saline (PBS) buffer (MilliporeSigma) and plated on LB agar plates with selective antibiotics. To count transformed *B. subtilis*, cells were plated directly on LB agar plates with selective antibiotics without dilution. LB agar plates were incubated at 37 °C with shaking (250 rpm) overnight and colonies were counted the next day. Transformation frequency was defined as the ratio of transformed *B. subtilis* to the total *B. subtilis* with detection limit ∼ 10^−9^. HGT efficiency was defined as the transformation frequency in co-culture with the *E. coli* donor. For DNase treatment, 1 unit/mL DNase I (Thermo Fisher Scientific) was added to the cell culture. One unit of DNase I completely degrades 1 μg of plasmid DNA in 10 min at 37 °C according to the manufacturer’s specification. To induce the lysis of *E. coli* in co-culture, 0, 0.02, 0.2, and 2 mM IPTG was added at the onset of the co-culture to induce the lysis gene *E* on pYC01 plasmid. To heat-kill *E. coli*, overnight *E. coli* cell culture was diluted to OD0.1 in LB and cultured at 37 °C with shaking (250 rpm) for 6h. Then the cells were incubated at 60 °C with shaking (250 rpm) for 30 min. Heat-killed cells were spun down at 5000g for 5 min. Pellet was resuspended with fresh LB and stored at 4 °C for use in the HGT experiment on the next day. CFU counting was performed after heat-killing to make sure that *E. coli* was completely killed. To inhibit *E. coli* growth in co-culture, 5 μg/mL chloramphenicol was at the onset of the co-culture. *B. subtilis* was not affected by chloramphenicol because of the introduced chloramphenicol acetyltransferase *cat* gene. Antibiotics used in this study were spectinomycin (spec) from Dot Scientific, chloramphenicol (chlor) from MilliporeSigma, MLS (1 μg/mL erythromycin from MilliporeSigma and 25 μg/mL lincomycin from Thermo Fisher Scientific), carbenicillin (carb) from MilliporeSigma, streptomycin (strep) from MilliporeSigma, and kanamycin (kan) from MilliporeSigma.

### qPCR measurements of extracellular DNA

To measure eDNA concentration, filtered supernatant from HGT experiments was added to a qPCR reaction mix containing 2X SsoAdvanced™ Universal Probes Supermix (Bio-Rad) and PrimeTime qPCR Assays (Integrated DNA Technologies) with 500 nM of each primer and 250 nM of probe. Each biological replicate had three technical replicates on the 96-well PCR plate (Thermo Fisher Scientific). qPCR primers can amplify *specR* on the pBB275 plasmid, *ampR* on the pUC19 plasmid, *ermR* introduced into *E. coli* MG1655 gDNA donor genome, and *caiE* in the *E. coli* MG1655 plasmid donor genome. qPCR primer sequences are listed in **Supplementary Table 4**. DNA standards of plasmid or gDNA were included in the qPCR run to determine the eDNA concentration of samples. DNA standards were previously quantified using Quant-iT dsDNA Assay Kit (Thermo Fisher Scientific). qPCR reactions were performed on the CFX Connect Real-Time PCR Detection System (Bio-Rad). eDNA concentration was calculated from the standard curve.

### Agarose gel electrophoresis for plasmid imaging

To determine the size of plasmids, 0.8% agarose (MIDSCI) gel was ran at 120V for ∼40 minutes. Before electrophoresis, extracted plasmids were mixed with Purple Gel Loading Dye (6X, no SDS) (New England Biolabs) and loaded into wells in the agarose gel. Quick-Load® Purple 1 kb DNA Ladder (New England Biolabs) or Quick-Load® 1 kb Extend DNA Ladder (New England Biolabs) was used for the reference of DNA length. Gel images were inverted by ImageJ for better visualization.

### Plate reader measurements of bacterial growth

To measure the growth rate of *recA*+ and *recA*-*E. coli* donors, cells from overnight culture were diluted to OD0.1 in LB and transferred to 96-well black and clear-bottom CELLSTAR® format sterile cell culture microplates (Greiner Bio-One) with 3 technical replicates. The plate was sealed with Breathe-Easy Adhesive Microplate Seal (Thermo Fisher Scientific) and cultured in the TECAN Spark 10M Multimode Microplate Reader for time-series OD measurements with shaking. A delay growth model was fitted to the growth curve to infer the cell doubling time (**Supplementary Information**).

### *Flow cytometer measurements of SOS response* in *E. coli* population

To characterize SOS response in *E. coli* population, cell culture of different *E. coli* strains with the SOS response reporter plasmid pBbA6c_PsulA_sfGFP was diluted to OD0.1 and cultured at 37 °C with shaking (250 rpm) for 6h. GFP expression was measured by the LSRFortessa X-20 Flow Cytometer (BD Biosciences). A blue (488 nm) laser was used for GFP excitation and a 530/30 nm filter was used for GFP emission. To quantify the percentage of SOS response activated cells, a threshold of GFP 10000 (a.u.) was used to determine GFP-ON cells. Data analysis was performed using a customized MATLAB script. The distribution of GFP and Forward Scatter was plotted by FlowJo.

### Microscopic imaging of single cells

To image single cells of *E. coli* and *B. subtilis* in the co-culture, overnight culture of each strain was first diluted to OD0.1 into LB plus 50 mM xylose. Cells were co-cultured for 3 h and 4 μL cell culture was transferred to the glass slide. 5 μL 0.1 % (w/v) Poly-L-lysine (Millipore Sigma) was spread evenly on the glass slide to help fix the cells. Pipetting was avoided to prevent the disruption of the physically associated cells. Single cells were imaged by Ti-E Eclipse inverted microscope (Nikon) with 40X magnification. GFP, RFP, and phase-contrast images were taken from multiple spots on the glass slide. Filters used for GFP (Chroma) were 470 nm/40 nm (excitation) and 525/50 nm (emission). Filters used for RFP (Chroma) were 560 nm/40 nm (excitation) and 630/70 nm (emission). To image SOS response of single cells, cell culture of *E. coli* MG1655 pBB275 or pUC19 with pBbA6c_PsulA_sfGFP plasmid was diluted to OD0.1 and cultured at 37 °C with shaking (250 rpm) for 6h. 4 μL cell culture was transferred to the glass slide and imaged by Nikon Eclipse Ti Microscope with 40X magnification. Phase-contrast and GFP images were taken to determine the presence of filamentous and fluorescent cells in *E. coli* MG1655 pBB275.

### Bioinformatic analysis of plasmid transfer to Bacillus and non-Bacillus bacteria

To search *E. coli* plasmid replication origins in the Genus *Bacillus*, ColE1 (589 bp in pUC19, Addgene Plasmid #50005), p15A (712 bp in pBbA6c-RFP, Addgene Plasmid #35290), CloDF13 (739 bp in pCSaE500, Addgene Plasmid #53182), and pSC101 (2224 bp in pBbS2k-RFP, Addgene Plasmid #35330) were searched in NCBI RefSeq Genome Database. Nucleotide BLAST was optimized for highly similar sequences. The organism was limited to *Bacillus* (taxid:1386). Max target sequences 5000 were selected. To search *E. coli* plasmid replication origins in non-*Bacillus* bacteria, the organism was limited to Bacteria (taxid:2) and excluded *Escherichia* (taxid:561) and *Bacillus* (taxid:1386). Accession date of data is 2021-09-03. Hits were listed in **Supplementary Table 1**. Hits with > 90% identity and > 80% percentage were selected for the analysis of sequence length. Sequence length longer than 10^6^ bp was defined as genomic integration. Shorter sequence length may be due to the contamination of plasmid DNA during sequencing steps^64^. Phylogenetic tree was constructed by aligning 16S rRNA sequences using Neighbor Joining method in MEGA v10.2.6^65^. To annotate genes near plasmid origin replication in *Bacillus* genomes, GenBank sequences were downloaded from NCBI website and plotted using SnapGene Viewer v5.3.2.

## ACKNOWLEDGEMENTS

We would like to thank Zhengyi Chen for the assistance with experiments and Michael Cox for sharing *E. coli* RecA strains and helpful suggestions. This work was supported by the Defense Advanced Research Projects Agency (DARPA) under grant number HR0011-19-2-0002 and National Institutes of Health under grant number R35GM124774.

## AUTHOR CONTRIBUTIONS

Y.Y.C., B.M.B. and O.S.V. conceived the research. Y.Y.C. performed the experiments. Y.Y.C. and Z.Z. performed bioinformatic analyses. J.M.P. constructed the *E. coli* strains. J.D.Z. constructed the pBB275 plasmids. T.G.F. assisted with the strain construction. K. A. assisted with the bioinformatic analysis. Y.Y.C. and O.S.V. wrote the manuscript. O.S.V. secured the funding.

## REFERENCES

1. Soucy, S. M., Huang, J. & Gogarten, J. P. Horizontal gene transfer: Building the web of life. Nature Reviews Genetics 16, 472–482 (2015).

2. Ropars, J. et al. Adaptive horizontal gene transfers between multiple cheese-associated fungi. Curr. Biol. 25, 2562–2569 (2015).

3. Clarke, M., Maddera, L., Harris, R. L. & Silverman, P. M. F-pili dynamics by live-cell imaging. Proc. Natl. Acad. Sci. U. S. A. 105, 17978–17981 (2008).

4. Overballe-Petersen, S. et al. Bacterial natural transformation by highly fragmented and damaged DNA. Proc. Natl. Acad. Sci. U. S. A. 110, 19860–19865 (2013).

5. Johnston, C., Martin, B., Fichant, G., Polard, P. & Claverys, J.-P. Bacterial transformation: distribution, shared mechanisms and divergent control. Nat. Rev. Microbiol. 12, 181–196 (2014).

6. Lerminiaux, N. A. & Cameron, A. D. S. Horizontal transfer of antibiotic resistance genes in clinical environments. Can. J. Microbiol. 65, 34–44 (2019).

7. Jensen, A., Valdórsson, O., Frimodt-Møller, N., Hollingshead, S. & Kilian, M. Commensal Streptococci serve as a reservoir for β-lactam resistance genes in Streptococcus pneumoniae. Antimicrob. Agents Chemother. 59, 3529–3540 (2015).

8. Sauerbier, J., Maurer, P., Rieger, M. & Hakenbeck, R. Streptococcus pneumoniae R6 interspecies transformation: genetic analysis of penicillin resistance determinants and genome-wide recombination events. Mol. Microbiol. 86, 692–706 (2012).

9. Hakenbeck, R., Grebe, T., Zähner, D. & Stock, J. B. β-lactam resistance in Streptococcus pneumoniae: penicillin-binding proteins and non-penicillin-binding proteins. Mol. Microbiol. 33, 673–678 (1999).

10. Nagler, M., Insam, H., Pietramellara, G. & Ascher-Jenull, J. Extracellular DNA in natural environments: features, relevance and applications. Applied Microbiology and Biotechnology 102, (2018).

11. Ibáñez de Aldecoa, A. L., Zafra, O. & González-Pastor, J. E. Mechanisms and regulation of extracellular DNA release and its biological roles in microbial communities. Front. Microbiol. 8, 1390 (2017).

12. Whitchurch, C. B., Tolker-Nielsen, T., Ragas, P. C. & Mattick, J. S. Extracellular DNA required for bacterial biofilm formation. Science (80-.). (2002). doi:10.1126/science.295.5559.1487

13. Prudhomme, M., Attaiech, L., Sanchez, G., Martin, B. & Claverys, J.-P. Antibiotic stress induces genetic transformability in the human pathogen streptoccus pneumoniae. Science (80-.). 313, 89–92 (2006).

14. Finkel, S. E. & Kolter, R. DNA as a nutrient: Novel role for bacterial competence gene homologs. J. Bacteriol. (2001). doi:10.1128/JB.183.21.6288-6293.2001

15. Zafra, O., Lamprecht-Grandío, M., de Figueras, C. G. & González-Pastor, J. E. Extracellular DNA Release by Undomesticated Bacillus subtilis Is Regulated by Early Competence. PLoS One (2012). doi:10.1371/journal.pone.0048716

16. Boman, H. G. & Eriksson, K. G. Penicillin-induced lysis in Escherichia coli. J. Gen. Microbiol. 31, (1963).

17. Islam, M. S., Aryasomayajula, A. & Selvaganapathy, P. R. A review on macroscale and microscale cell lysis methods. Micromachines 8, (2017).

18. Cooper, R. M., Tsimring, L. & Hasty, J. Inter-species population dynamics enhance microbial horizontal gene transfer and spread of antibiotic resistance. Elife 6, e25950 (2017).

19. Borgeaud, S., Metzger, L. C., Scrignari, T. & Blokesch, M. The type VI secretion system of Vibrio cholerae fosters horizontal gene transfer. Science (80-.). 347, 63–67 (2015).

20. Wholey, W.-Y., Kochan, T. J., Storck, D. N. & Dawid, S. Coordinated bacteriocin expression and competence in Streptococcus pneumoniae contributes to genetic adaptation through neighbor predation. PLoS Pathog. 12, e1005413 (2016).

21. Zhang, X. et al. Stress-induced, highly efficient, donor cell-dependent cell-to-cell natural transformation in Bacillus subtilis. J. Bacteriol. 200, e00267–18 (2018).

22. Tribble, G. D. et al. Natural competence is a major mechanism for horizontal DNA transfer in the oral pathogen Porphyromonas gingivalis. MBio 3, e00231–11 (2012).

23. Stewart, G. J., Carlson, C. A. & Ingraham, J. L. Evidence for an active role of donor cells in natural transformation of Pseudomonas stutzeri. J. Bacteriol. 156, 30–35 (1983).

24. Paul, J. H., Thurmond, J. M., Frischer, M. E. & Cannon, J. P. Intergeneric natural plasmid transformation between E. coli and a marine Vibrio species. Mol. Ecol. 1, 37–46 (1992).

25. Pavlopoulou, A. RecA: A universal drug target in pathogenic bacteria. Front. Biosci. - Landmark 23, (2018).

26. Kowalczykowski, S. C. & Eggleston, A. K. Homologous pairing and DNA strand-exchange proteins. Annual Review of Biochemistry 63, (1994).

27. Johnson, A. D. et al. λ Repressor and cro - Components of an efficient molecular switch. Nature 294, (1981).

28. Beaber, J. W., Hochhut, B. & Waldor, M. K. SOS response promotes horizontal dissemination of antibiotic resistance genes. Nature (2004). doi:10.1038/nature02241

29. Maslowska, K. H., Makiela-Dzbenska, K. & Fijalkowska, I. J. The SOS system: A complex and tightly regulated response to DNA damage. Environmental and Molecular Mutagenesis 60, (2019).

30. Simmons, L. A., Foti, J. J., Cohen, S. E. & Walker, G. C. The SOS Regulatory Network. EcoSal Plus 3, (2008).

31. Bedbrook, J. R. & Ausubel, F. M. Recombination between bacterial plasmids leading to the formation of plasmid multimers. Cell 9, (1976).

32. Canosi, U., Morelli, G. & Trautner, T. A. The relationship between molecular structure and transformation efficiency of some S. aureus plasmids isolated from B. subtilis. MGG Mol. Gen. Genet. 166, (1978).

33. Lopatkin, A. J. et al. Antibiotics as a selective driver for conjugation dynamics. Nat. Microbiol. (2016). doi:10.1038/nmicrobiol.2016.44

34. Zhang, X.-Z. & Zhang, Y.-H. P. Simple, fast and high-efficiency transformation system for directed evolution of cellulase in Bacillus subtilis. Microb. Biotechnol. 4, 98–105 (2011).

35. Hahn, J., Inamine, G., Kozlov, Y. & Dubnau, D. Characterization of comE, a late competence operon of Bacillus subtilis required for the binding and uptake of transforming DNA. Mol. Microbiol. 10, (1993).

36. Blattner, F. R. et al. The complete genome sequence of Escherichia coli K-12. Science 277, (1997).

37. Grant, S. G. N., Jessee, J., Bloom, F. R. & Hanahan, D. Differential plasmid rescue from transgenic mouse DNAs into Escherichia coli methylation-restriction mutants. Proc. Natl. Acad. Sci. U. S. A. 87, (1990).

38. Erental, A., Kalderon, Z., Saada, A., Smith, Y. & Engelberg-Kulka, H. Apoptosis-Like Death, An extreme SOS response in Escherichia Coli. MBio 5, (2014).

39. McCool, J. D. et al. Measurement of SOS expression in individual Escherichia coli K-12 cells using fluorescence microscopy. Mol. Microbiol. 53, (2004).

40. Robinson, A. et al. Regulation of Mutagenic DNA Polymerase V Activation in Space and Time. PLoS Genet. 11, (2015).

41. Arís, A. et al. The expression of recombinant genes from bacteriophage lambda strong promoters triggers the SOS response in Escherichia coli. Biotechnol. Bioeng. 60, (1998).

42. Lee, J. et al. Interplay of SOS induction, recombinant gene expression, and multimerization of plasmid vectors in Escherichia coli. Biotechnol. Bioeng. 80, (2002).

43. Johnson, S. A. et al. Increasing the bactofection capacity of a mammalian expression vector by removal of the f1 ori. Cancer Gene Ther. 26, (2019).

44. Zeigler, D. R. et al. The origins of 168, W23, and other Bacillus subtilis legacy strains. J. Bacteriol. 190, (2008).

45. Goto, A. & Kunioka, M. Biosynthesis and Hydrolysis of Poly(γ-glutamic acid) from Bacillus subtilis IF03335. Biosci. Biotechnol. Biochem. 56, (1992).

46. Wang, P. et al. Development of an efficient conjugation-based genetic manipulation system for Pseudoalteromonas. Microb. Cell Fact. 14, (2015).

47. García-Bayona, L. & Comstock, L. E. Bacterial antagonism in host-associated microbial communities. Science 361, (2018).

48. Maurice, C. F., Haiser, H. J. & Turnbaugh, P. J. Xenobiotics shape the physiology and gene expression of the active human gut microbiome. Cell 152, (2013).

49. Usajewicz, I. & Nalepa, B. Survival of Escherichia coli O157:H7 in milk exposed to high temperatures and high pressure. Food Technol. Biotechnol. 44, (2006).

50. Frenkel, L. & Bremer, H. Increased Amplification of Plasmids pBR322 and pBR327 by Low Concentrations of Chloramphenicol. DNA 5, (1986).

51. Berry, J. L., Cehovin, A., McDowell, M. A., Lea, S. M. & Pelicic, V. Functional Analysis of the Interdependence between DNA Uptake Sequence and Its Cognate ComP Receptor during Natural Transformation in Neisseria Species. PLoS Genet. 9, (2013).

52. Haudiquet, M., Buffet, A., Rendueles, O. & Rocha, E. P. C. Interplay between the cell envelope and mobile genetic elements shapes gene flow in populations of the nosocomial pathogen klebsiella pneumoniae. PLoS Biol. 19, (2021).

53. Lopatkin, A. J. et al. Antibiotics as a selective driver for conjugation dynamics. Nat. Microbiol. (2016). doi:10.1038/nmicrobiol.2016.44

54. Wein, T., Hülter, N. F., Mizrahi, I. & Dagan, T. Emergence of plasmid stability under non-selective conditions maintains antibiotic resistance. Nat. Commun. 10, (2019).

55. Dubey, G. P. & Ben-Yehuda, S. Intercellular nanotubes mediate bacterial communication. Cell 144, 590–600 (2011).

56. Bhattacharya, S. et al. A Ubiquitous Platform for Bacterial Nanotube Biogenesis. Cell Rep. 27, (2019).

57. Rahmer, R., Heravi, K. M. & Altenbuchner, J. Construction of a super-competent Bacillus subtilis 168 using the PmtlA-comKS inducible cassette. Front. Microbiol. 6, 1431 (2015).

58. Evans, D. R. et al. Systematic detection of horizontal gene transfer across genera among multidrug-resistant bacteria in a single hospital. Elife 9, (2020).

59. Ronda, C., Chen, S. P., Cabral, V., Yaung, S. J. & Wang, H. H. Metagenomic engineering of the mammalian gut microbiome in situ. Nat. Methods (2019). doi:10.1038/s41592-018-0301-y

60. Lutz, R. & Bujard, H. Independent and tight regulation of transcriptional units in escherichia coli via the LacR/O, the TetR/O and AraC/I1-I2 regulatory elements. Nucleic Acids Res. 25, 1203–1210 (1997).

61. Egbert, R. G. et al. A versatile platform strain for high-fidelity multiplex genome editing. Nucleic Acids Res. 47, 3244–3256 (2019).

62. Overkamp, W. et al. Benchmarking various green fluorescent protein variants in Bacillus subtilis, Streptococcus pneumoniae, and Lactococcus lactis for live cell imaging. Appl. Environ. Microbiol. 79, 6481–6490 (2013).

63. Konkol, M. A., Blair, K. M. & Kearns, D. B. Plasmid-encoded comi inhibits competence in the ancestral 3610 strain of Bacillus subtilis. J. Bacteriol. 195, (2013).

64. Wally, N. et al. Plasmid DNA contaminant in molecular reagents. Sci. Rep. 9, (2019).

65. Kumar, S., Stecher, G., Li, M., Knyaz, C. & Tamura, K. MEGA X: Molecular evolutionary genetics analysis across computing platforms. Mol. Biol. Evol. 35, (2018).

