## Supplementary Information for "Efficient plasmid transfer via natural competence in a synthetic microbial community"

**This file includes:**
Description of the delay growth model.
**Supplementary Figure 1**. Control experiments for the plasmid transfer in community.
**Supplementary Figure 2**. Characterization of gDNA transfer in community.
**Supplementary Figure 3**. Growth of *recA*+ and *recA*- *E. coli* donors.
**Supplementary Figure 4**. Plasmid maps for characterizing HGT efficiency, plasmid multimerization, and eDNA release.
**Supplementary Figure 5**. Induction of SOS response in *E. coli* MG1655 pBB275.
**Supplementary Figure 6**. Gel image of the plasmid in the supernatant.
**Supplementary Figure 7**. DNA release from the heat-killed, chloramphenicol-inhibited, and live *E. coli*.
**Supplementary Figure 8**. Nucleotide BLAST search results of ColE1, p15A, pSC101, and CloDF13 replication origins in *Bacillus* and non-*Bacillus* DNA sequences.
**Supplementary Figure 9**. Presence of ColE1 replication origin in *Bacillus* and non-*Bacillus* genomes.
**Supplementary Table 1.** Nucleotide BLAST search results of *E. coli* plasmid replication origins in *Bacillus* and non-*Bacillus* genome sequences.
**Supplementary Table 2**. List of plasmids.
**Supplementary Table 3**. List of bacterial strains.
**Supplementary Table 4**. Sequences of qPCR primers.

**Delay growth model for inferring doubling time from growth curve**

To estimate the doubling time of *E. coli* from the growth curve, a coupled ordinary differential equation model was fit to the time-series measurements of absorbance at 600 nm (OD)[1]. The two species $E_n$ and $E_g$ represent the sub-populations of non-growing and growing cells, respectively. The equations for this growth model are

$$\frac{dE_n}{dt} = -\,k_g E_n, \tag{S1}$$

$$\frac{dE_g}{dt} = \;k_g E_n + \mu_e E - \alpha_{ee} E^2\,, \tag{S2}$$

where $k_g$ represents the transition rate from the non-growing sub-population $E_n$ to growing sub-population $E_g$, $\mu_e$ denotes the exponential growth rate, and $\alpha_{ee}$ represents the intra-species interaction coefficient of the growing sub-population. We fit the model to the time-series OD measurements  using custom code in MATLAB[2]. A nonlinear programming solver was used to minimize the mean squared error between the model prediction and experimental measurement across all time points. The doubling time $t_D$ was computed using the equation $t_D$ = $\log_e 2/\mu_e$.

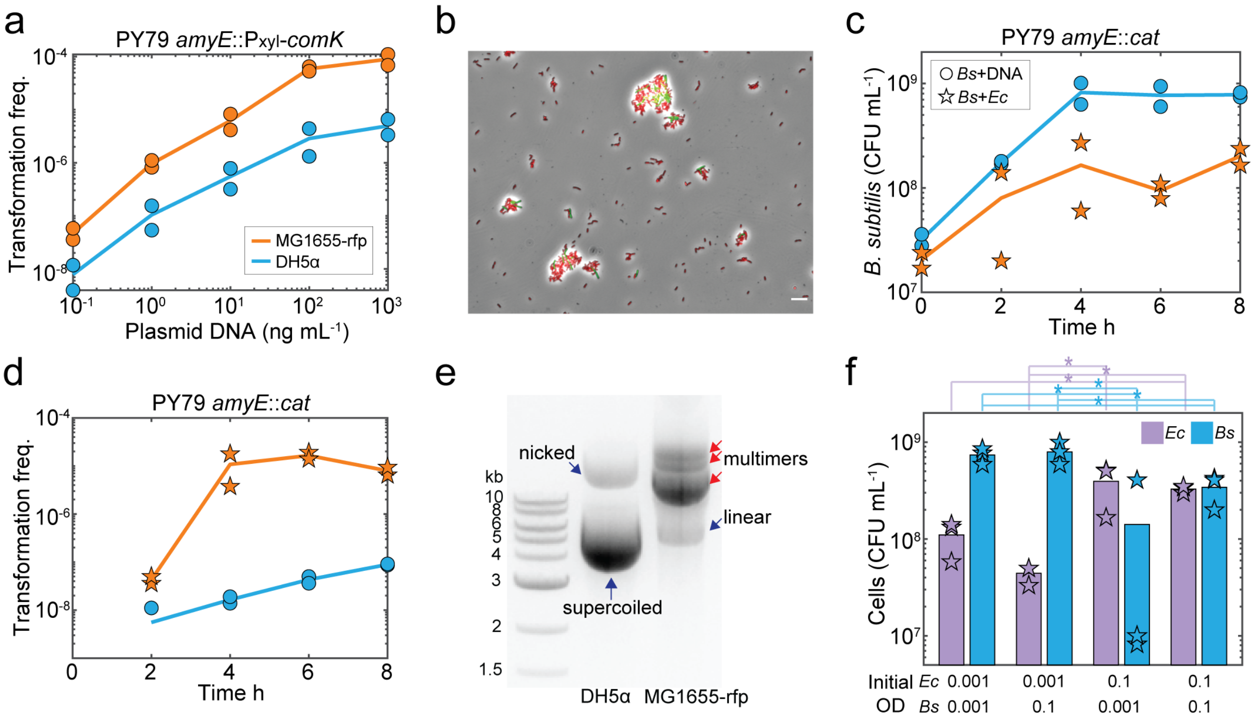


**Supplementary Figure 1. Characterization of plasmid transfer in the synthetic microbial community. (a)** Scatter plot of the initial concentration of pBB275 plasmid DNA introduced into a monoculture of xylose-inducible *comK B. subtilis* and transformation frequency at 6 hr. The plasmid derived from *E.coli* MG1655-rfp displayed ~10-fold higher transformation frequency than plasmid derived from *E. coli* DH5α. **(b)** Microscopy image of *E. coli* MG1655-rfp (red) and xylose-inducible *comK B. subtilis* (green) in the microbial community at 3 hr. Scale bar denotes 10 μm. **(c)** Time-series measurements of the abundance of wild-type *B. subtilis* in the monoculture and community with *E. coli* MG1655-rfp harboring pBB275. **(d)** Time-series measurements of transformation frequency of the pBB725 plasmid in the monoculture of wild-type *B. subtilis* and community with *E. coli* MG1655-rfp pBB275. In the monoculture, 100 ng/mL pBB275 plasmid DNA derived from *E. coli* DH5α was introduced. One biological replicate was below the detection limit of ~$10^{-9}$ transformation frequency at 2 hr. **(e)** Gel image of the integrative plasmid pBB275 extracted from *E. coli* MG1655-rfp or DH5α. Plasmids extracted from *E. coli* MG1655-rfp contained multimers indicated by red arrows. Multimers had larger molecular weight than monomers (nicked, linear, and supercoiled indicated by blue arrows) and migrated slower on the gel by electrophoresis. **(f)** Bar plot of the abundances of *E. coli* MG1655-rfp pBB275 or xylose-inducible *comK B. subtilis* 6 hr in the community inoculated with different initial species densities. Stars (*) denote unpaired *t*-test with *p*-values < 0.05. Low initial density of *E. coli* (OD0.001) yielded lower *E. coli* density and high *B. subtilis* at 6 hr, whereas high initial density of *E. coli* (OD0.1) yielded higher *E. coli* density and lower *B. subtilis* at 6 hr. Time-course and DNA titration experiments had two biological replicates. Initial cell density experiment had three biological replicates. Lines and bars are the average of the biological replicates.

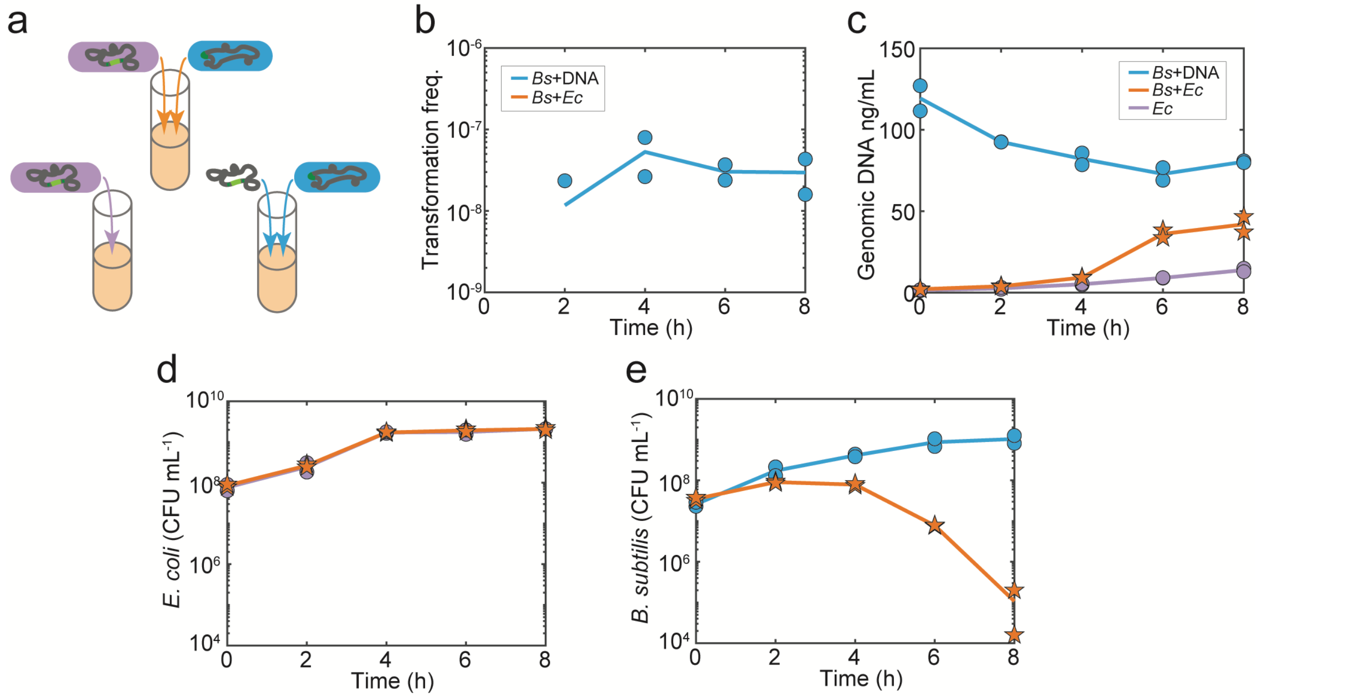


**Supplementary Figure 2. Characterization of horizontal gene transfer of genomic DNA in the microbial community. (a)** Schematic of experimental design to characterize the temporal changes in transfer of genomic DNA (gDNA) in the *B. subtilis* monoculture supplemented with 100 ng/mL purified *E. coli* MG1655 donor gDNA (blue) or community composed of the *E. coli* gDNA donor and xylose-inducible *comK B. subtilis* (orange). Extracellular gDNA release was quantified in the *E. coli* monoculture (purple). An erythromycin resistance gene flanked by two ~600 bp *B. subtilis* PY79 *yvbJ* was introduced onto the *E. coli* MG1655 genome. Time-series measurements of **(b)** the transformation frequency of *E. coli* MG1655 donor gDNA in monoculture or in the community, **(c)** extracellular *E. coli* MG1655 donor gDNA concentration, **(d)** abundance of the *E. coli* MG1655 gDNA donor, and **(e)** abundance of the xylose-inducible *comK B. subtilis* in the monoculture or community. One biological replicate in the *B. subtilis* monoculture was below detection limit of ~$10^{-9}$ transformation frequency at 2 hr. Transformation frequency of gDNA in the community was below the detection limit of ~$10^{-9}$. Each experiment had two biological replicates. Lines are the average of the biological replicates.

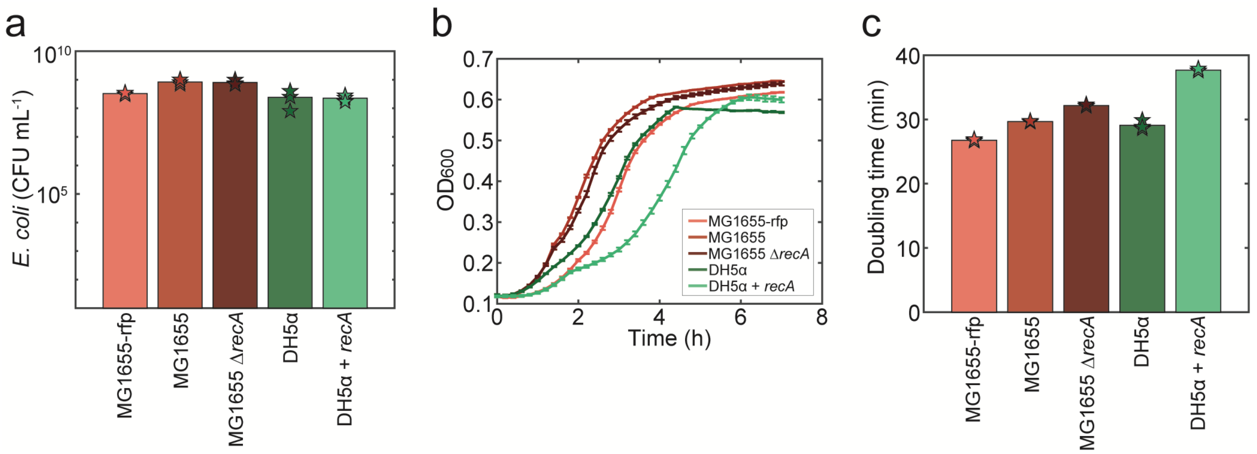


**Supplementary Figure 3. Growth characterization of *recA*+ or *recA*- *E. coli* donors. (a)** Abundance of the *recA*+ or *recA*- *E. coli* plasmid donors at 6 hr in community with xylose-inducible *comK B. subtilis*. All *E. coli* strains harbored the pBB275 plasmid. **(b)** Time-series measurements of absorbance at 600 nm ($OD_{600}$) for the *recA*+ or *recA*- *E. coli* plasmid donors in monoculture. **(c)** Inferred doubling times based on the growth responses of the *recA*+ or *recA*- *E. coli* plasmid donors using the delayed differential equation growth model. *E. coli* DH5α + *recA* displayed the slowest growth rate. Bars and lines are the average of three biological replicates. Error bars in **(b)** denote 1 s.d. from the mean.

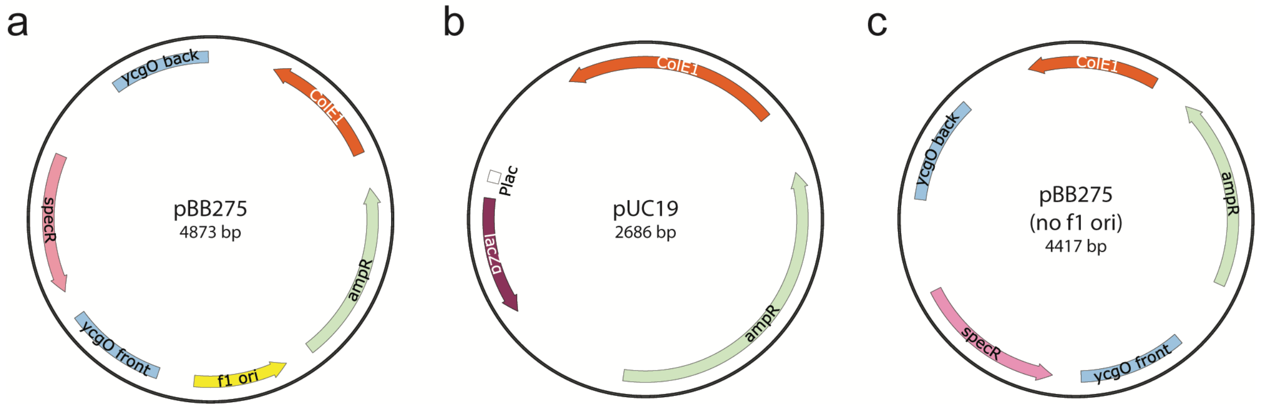


**Supplementary Figure 4. Plasmid maps for characterizing HGT efficiency, plasmid multimerization, and eDNA release in the synthetic community. (a)** Map of the pBB275 plasmid. *specR* denotes the spectinomycin resistance gene used for the selection of *B. subtilis* genomic integration in *ycgO* locus. The f1 ori denotes the phage replication origin and is a remnant of the original plasmid with the ColE1 replication origin and ampicillin resistance gene *ampR*. **(b)** Map of the pUC19 plasmid. pUC19 plasmid contains the same ColE1 replication origin and *ampR* as the pBB275 plasmid but does not contain the f1 ori, *ycgO*, and *specR*. This plasmid contains the β-galactosidase gene *lacZα* from *E. coli lac* operon. **(c)** Map of the pBB275 plasmid lacking the f1 origin.

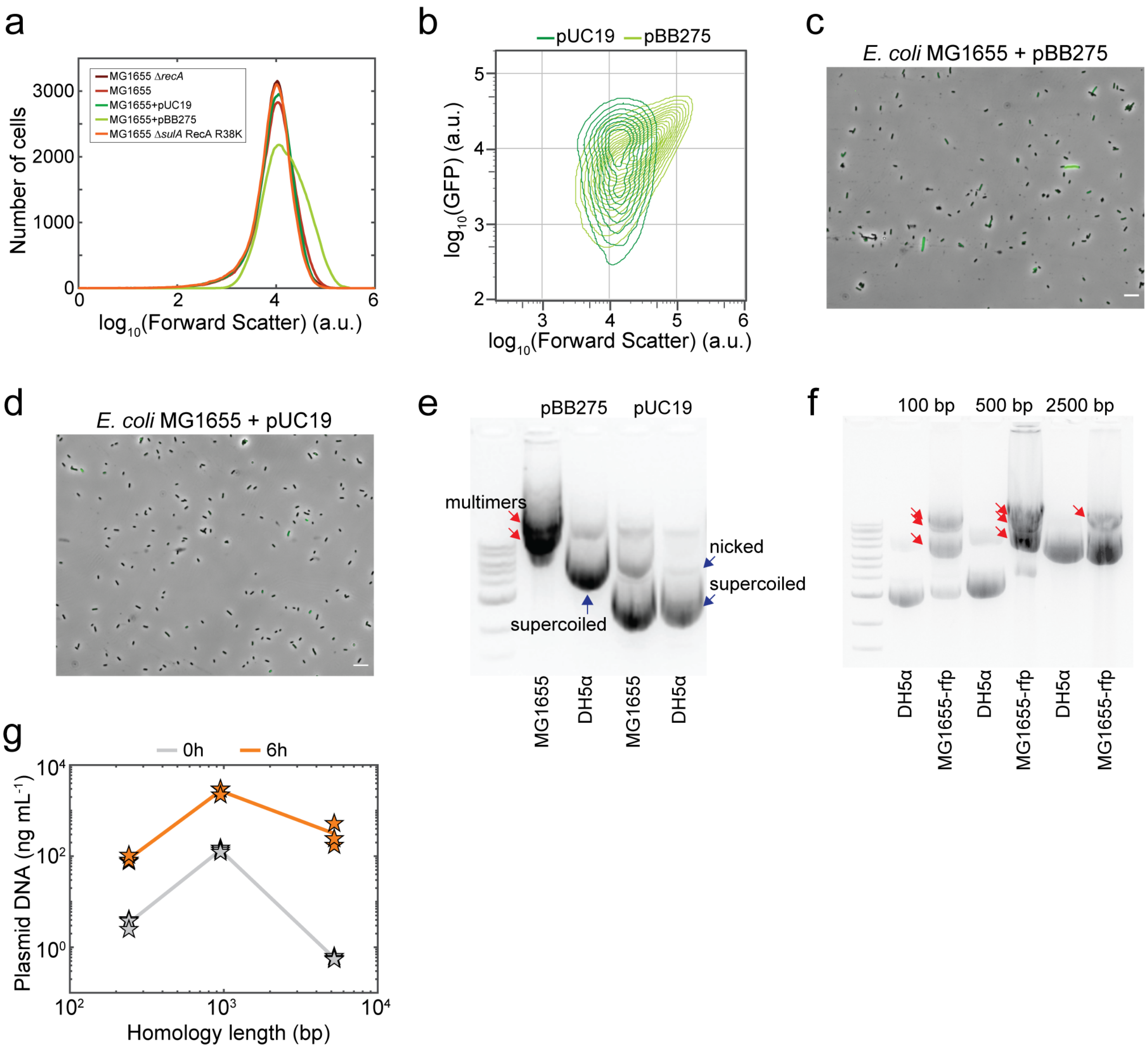


**Supplementary Figure 5. Induction of the SOS response in *E. coli* MG1655 harboring pBB275. (a)** Histogram of forward scatter based on flow cytometry of *E. coli* MG1655 Δ*recA*, *E. coli* MG1655, *E. coli* MG1655 pUC19, *E. coli* MG1655 pBB275, or *E. coli* MG1655 Δ*sulA* RecA E38K at 6 hr in monoculture. All *E. coli* strains harbored the SOS response promoter fusion to GFP on a plasmid. **(b)** Contour plot of GFP expression and forward scatter of *E. coli* MG1655 pUC19 or *E. coli* MG1655 pBB275 at 6 hr in monoculture based on flow cytometry. **(c)** Microscopy image of *E. coli* MG1655 harboring pBB275 and the SOS response reporter plasmid. Scale bar represents 10 μm. **(d)** Microscopy image of *E. coli* MG1655 harboring pUC19 and the SOS response reporter plasmid. Scale bar denotes 10 μm. **(e)** Gel image of pBB275 and pUC19 extracted from *E. coli* MG1655 and DH5α. A 1 kb DNA Ladder was used as a reference. pUC19 only contained monomers while pBB275 extracted from *E. coli* MG1655 contained multimers, based on the changes in the bands derived from *E. coli* MG1655 (*recA*+) and DH5α (*recA*-). **(f)** Gel image of the pBB275 plasmid with ~100 bp, ~500 bp, or ~2500 bp homology arms. All plasmids extracted from *E. coli* MG1655-rfp contained multimers whereas plasmids extracted from DH5α did not display multimers. A 1 kb DNA Ladder was used for reference. **(g)** Extracellular pBB275 plasmid concentration at 0 hr or 6 hr in community conmposed of *E. coli* MG1655-rfp pBB275 and xylose-inducible *comK B. subtilis*. The plasmids were constructed with three different total homology lengths: ~200 bp, ~1000 bp, or ~5000 bp. Each experiment had three biological replicates. Lines are the average of the biological replicates.

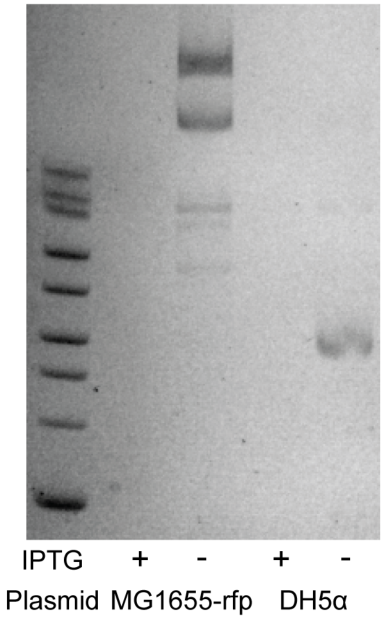


**Supplementary Figure 6. Gel image of pBB275 plasmid in the community supernatant.** The pBB275 plasmid was extracted from *E. coli* MG1655-rfp or DH5α and 1 µg/mL pBB275 plasmid DNA was introduced into the community composed of xylose-inducible *comK B. subtilis* and *E. coli* harboring an IPTG-inducible lysis gene. The supernatant of community was collected and filtered at 6 hr and 50 µL was measured by gel electrophoresis. These data show no band in the presence of 0.2 mM IPTG and multimeric or monomeric pBB275 were present in the absence of IPTG. A 1 kb Extend DNA Ladder was used as a reference.

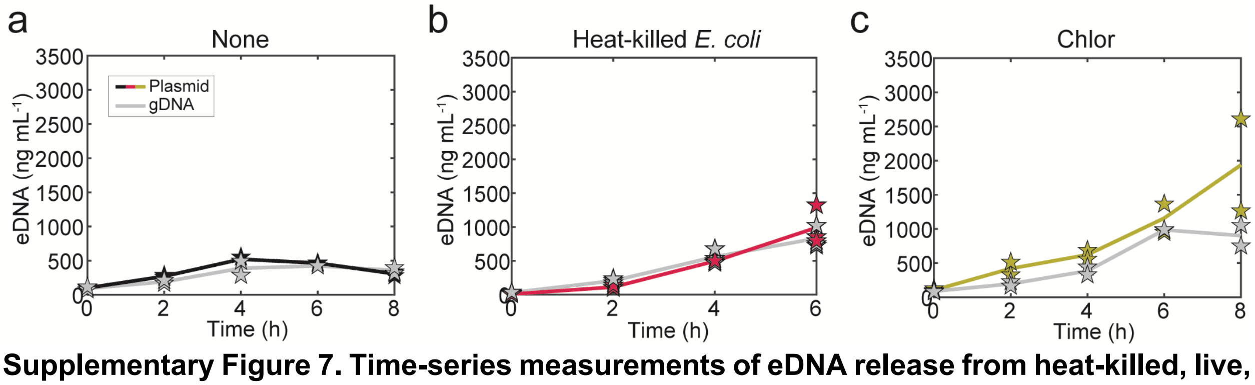


**Supplementary Figure 7. Time-series measurements of eDNA release from heat-killed, live, or chloramphenicol treated *E. coli*.** Time-series measurements of extracellular pBB275 plasmid or gDNA concentrations of the **(a)** live (not heat treated), **(b)** heat-killed, or **(c)** 5 µg/mL chloramphenicol treated *E. coli* MG1655-rfp pBB275 in a community with xylose-inducible *comK B. subtilis*. The heat-killed *E. coli* conditions had three biological replicates. Live and chloramphenicol treated *E. coli* experiments had two biological replicates. Lines are the average of the biological replicates. Note that the plasmid concentrations in **Supplementary Fig. 7a,c** are the same as **Fig. 4g** to compare with extracellular gDNA concentrations.

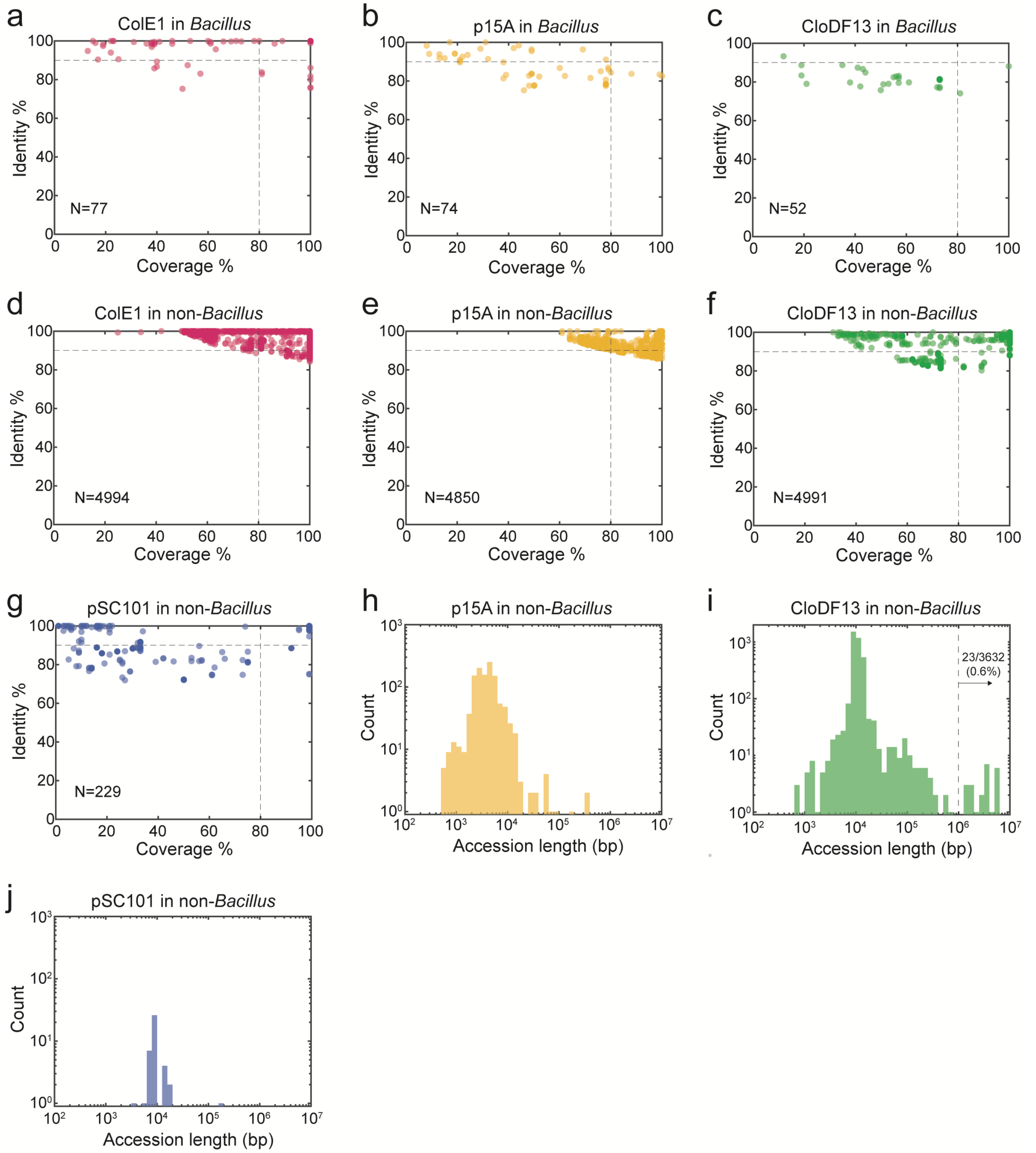


**Supplementary Figure 8. Nucleotide BLAST search results to identify ColE1, p15A, pSC101, and CloDF13 replication origins in *Bacillus* and non-*Bacillus* DNA sequences. (a)** Scatter plot of query coverage and percent identity of ColE1 in 77 *Bacillus* DNA sequences in the NCBI RefSeq Genome Database. Hits with >90% identity and >80% coverage were selected for the analysis of sequence length. Query coverage is the percentage of the hit sequence aligned to the query sequence. Percent identity is the percentage of nucleotides that match the alignment. **(b)**

Scatter plot of query coverage and percent identity of p15A in 74 *Bacillus* DNA sequences. **(c)** Scatter plot of query coverage and percent identity of CloDF13 in 52 *Bacillus* DNA sequences. **(d)** Scatter plot of query coverage and percent identity of ColE1 in 4994 non-*Bacillus* DNA sequences. **(e)** Scatter plot of query coverage and percent identity of p15A in 4850 non-*Bacillus* DNA sequences. **(f)** Scatter plot of query coverage and percent identity of CloDF13 in 4991 non-*Bacillus* DNA sequences. **(g)** Scatter plot of query coverage and percent identity of pSC101 in 229 non-*Bacillus* DNA sequences. Histograms of the sequence lengths of non-*Bacillus* DNA sequences with **(h)** p15A, **(i)** CloDF13, or **(j)** pSC101 replication origin.

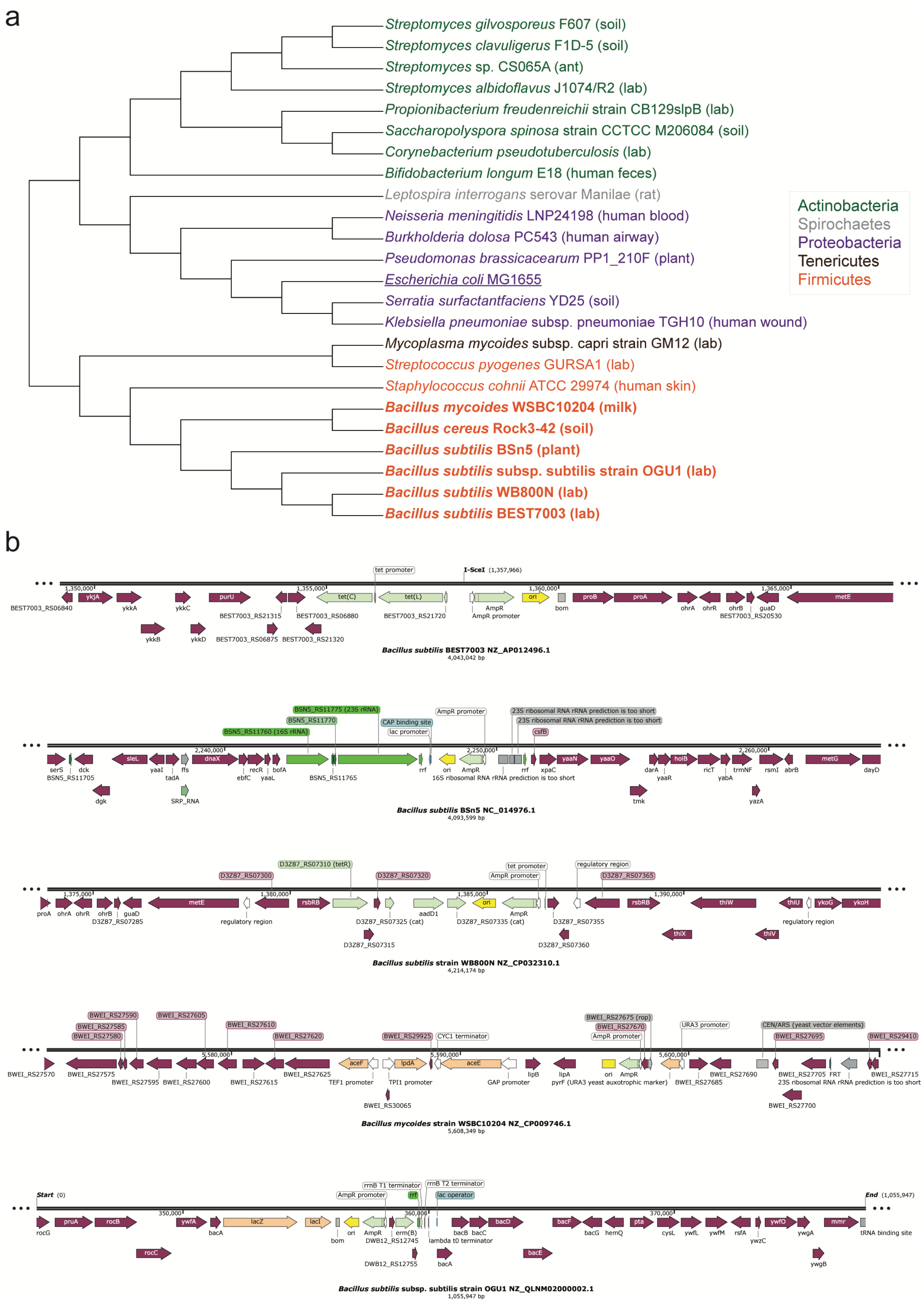
a
Streptomyces gilvosporeus F607 (soil)
Streptomyces clavuligerus F1D-5 (soil)
Streptomyces sp. CS065A (ant)
Streptomyces albidoflavus J1074/R2 (lab)
Propionibacterium freudenreichii strain CB129slpB (lab)
Saccharopolyspora spinosa strain CCTCC M206084 (soil)
Corynebacterium pseudotuberculosis (lab)
Bifidobacterium longum E18 (human feces)
Leptospira interrogans serovar Manilae (rat)
Neisseria meningitidis LNP24198 (human blood)
Burkholderia dolosa PC543 (human airway)
Pseudomonas brassicacearum PP1_210F (plant)
Escherichia coli MG1655
Serratia surfactantfaciens YD25 (soil)
Klebsiella pneumoniae subsp. pneumoniae TGH10 (human wound)
Mycoplasma mycoides subsp. capri strain GM12 (lab)
Streptococcus pyogenes GURSA1 (lab)
Staphylococcus cohnii ATCC 29974 (human skin)
Bacillus mycoides WSBC10204 (milk)
Bacillus cereus Rock3-42 (soil)
Bacillus subtilis BSn5 (plant)
Bacillus subtilis subsp. subtilis strain OGU1 (lab)
Bacillus subtilis WB800N (lab)
Bacillus subtilis BEST7003 (lab)
Actinobacteria
Spirochaetes
Proteobacteria
Tenericutes
Firmicutes
b
Bacillus subtilis BEST7003 NZ_AP012496.1
4,043,042 bp
Bacillus subtilis BSn5 NC_014976.1
4,093,599 bp
Bacillus subtilis strain WB800N NZ_CP032310.1
4,214,174 bp
Bacillus mycoides strain WSBC10204 NZ_CP009746.1
5,608,349 bp
Bacillus subtilis subsp. subtilis strain OGU1 NZ_QLNM02000002.1
1,055,947 bp

**Supplementary Figure 9. Presence of ColE1 replication origin in *Bacillus* and non-*Bacillus* genomes. (a)** Phylogenetic tree of bacteria that had a ColE1 replication origin identified in their genome. *E. coli* was included to compare the evolutionary distance. Different phyla are labeled with different colors. Known environments from which the bacteria were isolated are indicated. **(b)** Genome map of ColE1 and the flanking genes for five representative *Bacillus* genomes. Antibiotic resistance genes are labeled as light green. Heterologous genes are labeled as light orange. *Bacillus cereus* Rock3-42 was not included due to the scaffold gaps near ColE1.

**Supplementary Table 2. List of plasmids.**

| Plasmid | Genotype | Description |
|---|---|---|
| pBB275_100bp | *ycgO*(150 bp)-*specR*-*ycgO*(92 bp), f1 ori, ColE1, *ampR* | Integrative plasmid for *B. subtilis* transformation with ~100 bp *ycgO* homology arms |
| pBB275_500bp | *ycgO*(475 bp)-*specR*-*ycgO*(477 bp), f1 ori, ColE1, *ampR* | Integrative plasmid for *B. subtilis* transformation with ~500 bp *ycgO* homology arms |
| pBB275_2500bp | *ycgO*(2636 bp)-*specR*-*ycgO*(2583 bp), f1 ori, ColE1, *ampR* | Integrative plasmid for *B. subtilis* transformation with ~2500 bp *ycgO* homology arms |
| pUC19 | *lacZα*, ColE1, *ampR* | ColE1 plasmid without f1 origin and *B. subtilis* homology arms |
| pBB275 (no f1 ori) | *ycgO*(475 bp)-*specR*-*ycgO*(477 bp), ColE1, *ampR* | Integrative plasmid for *B. subtilis* transformation with ~500 bp *ycgO* homology arms but no f1 replication origin |
| pBbA6k_J23100_recA | $P_{J23100}$-*recA*, p15A, *kanR* | Constitutively expressed *recA* |
| pBbA6c_PsulA_sfGFP | $P_{sulA}$-*sfgfp*, p15A, *cat* | SOS response reporter |
| pYC01 | $P_{A1lacO-1}$-*E*, CloDF13, *smR* | IPTG-inducible phage φX174 lysis gene *E* |

**Supplementary Table 3. List of bacterial strains.**

| Strain | Genotype | Description | Figures |
|---|---|---|---|
| msOSV01034 | *E. coli* MG1655 *caiE*::$P_{J23100}$-*mCherry* (pBB275_500bp) | *E. coli* MG1655-rfp plasmid donor with pBB275_500bp | 1, 2A-C, 2H, 3I, 4, S1A, S1C-F, S3, S5F-G, S6, and S7 |
| msOSV01033 | *B. subtilis* PY79 *amyE*::$P_{xylA}$-*comK*, *cat*; *lacA*::*ermR* | Xylose-inducible comK *B. subtilis* for plasmid transformation | 1, 2A-C, 2E-F, 2H, 3, 4, S1A, S1F, |

| | | | |
|---|---|---|---|
| | | | S5G, S6, and S7 |
| msOSV00858 | *B. subtilis* PY79 *amyE*:: *cat*; *lacA*::*ermR* | *B. subtilis* without xylose-inducible *comK* for plasmid transformation | S1C-D |
| msOSV00389 | *E. coli* DH5α (pBB275_500bp) | *E. coli* DH5α plasmid donor with pBB275_500bp | 1B-F, 2A-B, 2H, 3I, S1A, S1C-E, S3, S5E-F, and S6 |
| msOSV01032 | *B. subtilis* PY79 *comEC*::*kanR*; *amyE*::$P_{xylA}$-*comK*, *cat*; *lacA*::*ermR* | Xylose-inducible comK *B. subtilis* without *comEC* | 1F |
| msOSV00348 | *E. coli* MG1655 *caiE*::*yvbJ*(595 bp)-*ermR*-*yvbJ*(594 bp) | *E. coli* MG1655 gDNA donor | S2 |
| msOSV00184 | *B. subtilis* PY79 *amyE*::$P_{xylA}$-*comK*, *cat; yhdGH*::*kanR* | Xylose-inducible comK *B. subtilis* for *E. coli* gDNA transformation | S2 |
| msOSV01107 | *E. coli* MG1655 (pBB275_500bp) | *E. coli* MG1655 plasmid donor with pBB275_500bp | 2A-C, 2E-F, S3, and S5E |
| msOSV01105 | *E. coli* MG1655 Δ*recA* (pBB275_500bp) | *E. coli* MG1655 Δ*recA* plasmid donor with pBB275_500bp | 2A-C and S3 |
| msOSV01106 * | *E. coli* DH5α (pBB275_500bp, pBbA6k_J23100_recA) | *E. coli* DH5α+*recA* plasmid donor with pBB275_500bp and pBbA6k_J23100_recA | 2A-B and S3 |
| msOSV01115 | *E. coli* MG1655 Δ*recA* (pBbA6c_PsulA_sfGFP) | *E. coli* MG1655 Δ*recA* with SOS response reporter plasmid | 2D and S5A |
| msOSV01116 | *E. coli* MG1655 (pBbA6c_PsulA_sfGFP) | *E. coli* MG1655 with SOS response reporter plasmid | 2D and S5A |
| msOSV01141 | *E. coli* MG1655 (pBbA6c_PsulA_sfGFP, pUC19) | *E. coli* MG1655 with SOS response reporter plasmid and pUC19 plasmid | 2D, S5A-B, and S5D |
| msOSV01119 | *E. coli* MG1655 (pBbA6c_PsulA_sfGFP, pBB275_500bp) | *E. coli* MG1655 with SOS response reporter plasmid and pBB275_500bp plasmid | 2D and S5A-C |
| msOSV01124 | *E. coli* MG1655 Δ*sulA* RecA E38K::*kanR* (pBbA6c_PsulA_sfGFP) | *E. coli* MG1655 Δ*sulA* RecA E38K with SOS response reporter plasmid showing constitutive SOS response | 2D and S5A |
| msOSV01138 | *E. coli* MG1655 (pUC19) | *E. coli* MG1655 with pUC19 plasmid | 2E and S5E |
| msOSV01139 | *E. coli* DH5α (pUC19) | *E. coli* DH5α with pUC19 plasmid | S5E |
| msOSV01254 | *E. coli* MG1655 (pBB275 (no f1 ori)) | *E. coli* MG1655 plasmid donor with pBB275 (no f1 ori) plasmid | 2F |

| | | | |
|---|---|---|---|
| msOSV01036 | *E. coli* MG1655 *caiE*::$P_{J23100}$-*mCherry* (pBB275_100bp) | *E. coli* MG1655-rfp plasmid donor with pBB275_100bp | 2H and S5F-G |
| msOSV01037 | *E. coli* MG1655 *caiE*::$P_{J23100}$-*mCherry* (pBB275_2500bp) | *E. coli* MG1655-rfp plasmid donor with pBB275_2500bp | 2H and S5F-G |
| msOSV00387 | *E. coli* DH5α (pBB275_2500bp) | *E. coli* DH5α plasmid donor with pBB275_2500bp | 2H and S5F |
| msOSV00388 | *E. coli* DH5α (pBB275_100bp) | *E. coli* DH5α plasmid donor with pBB275_100bp | 2H and S5F |
| msOSV01050 | *E. coli* BW29427 (pBB275_100bp) | DAP-auxotrophic *E. coli* plasmid donor with pBB275_100bp | 2I |
| msOSV01051 | *E. coli* BW29427 (pBB275_500bp) | DAP-auxotrophic *E. coli* plasmid donor with pBB275_500bp | 2I |
| msOSV01030 | *E. coli* BW29427 (pBB275_2500bp) | DAP-auxotrophic *E. coli* plasmid donor with pBB275_2500bp | 2I |
| msOSV00150 | *B. subtilis* 168 | Wild-type *B. subtilis* 168 | 2I |
| msOSV00117 | *B. subtilis* natto IFO3335 | Wild-type *B. subtilis* natto IFO3335 | 2I |
| usOSV00230 | *B. subtilis* PY79 | Wild-type *B. subtilis* PY79 | 2I |
| msOSV00608 | *E. coli* MG1655 *caiE*::$P_{J23100}$-*mCherry* (pBB275_500bp, pYC01) | *E. coli* MG1655-rfp plasmid donor with pBB275_500bp and IPTG-inducible lysis plasmid | 3A-H |
| msOSV01027 | *E. coli* MG1655 *caiE*::$P_{J23100}$-*mCherry* (pYC01) | *E. coli* MG1655-rfp with IPTG-inducible lysis plasmid | 3I and S6 |
| msOSV00105 | *E. coli* MG1655 *caiE*::$P_{J21300}$-*mCherry* | RFP-labeled *E. coli* for single-cell imaging | 3I, S1B |
| msOSV00837 | *B. subtilis* PY79 *ycgO*::$P_{hyperspank}$-*gfp*(Sp), *cat*; *lacA*::$P_{xylA}$-*comK*, *ermR* | GFP-labeled *B. subtilis* for single-cell imaging | S1B |

* Kanamycin was not used for culturing *E. coli* since it may impact *B. subtilis* transformation in community.

**Supplementary Table 4. Sequences of qPCR primers.**

| Part | Sequence |
|---|---|
| pBB275(*specR*)_FW | CCCTATGTTCTAATGGAGAAGATTCA |
| pBB275(*specR*)_RV | ATCAGGATGATGAAACCAACTCT |
| pBB275(*specR*)_Probe | /56-FAM/AGATATTGC/ZEN/GGGAAATGCAGTGGC/3IABkFQ/ |
| pUC19(*ampR*)_FW | CCCAACTGATCTTCAGCATCTT |
| pUC19(*ampR*)_RV | TTTCCGTGTCGCCCTTATTC |
| pUC19(*ampR*)_Probe | /56-FAM/ACTTTCACC/Zen/AGCGTTTCTGGGTGA/3IABkFQ/ |
| *E. coli* MG1655(*caiE*::*ermR*)_FW | GGTTGATAATGAACTGTGCTGAT |

| | |
|---|---|
| *E. coli* MG1655(*caiE*::*ermR*)_RV | CGCATCCGATTGCAGTATAAAT |
| *E. coli* MG1655(*caiE*::*ermR*)_Probe | /56-FAM/CATCATGTT/ZEN/CATATTTATCAGAGCTCGTGC/3IABkFQ/ |
| *E. coli* MG1655(*caiE*)_FW | ATACGCGGCGTCTTTCTTAC |
| *E. coli* MG1655(*caiE*)_RV | AGTGTCAGCGACTGGTTAAAG |
| *E. coli* MG1655(*caiE*)_Probe | /56-FAM/ATCGATGAG/ZEN/CAAATTTACGCGCGC/3IABkFQ/ |

**REFERENCES**

1. Fridman, O., Goldberg, A., Ronin, I., Shoresh, N. & Balaban, N. Q. Optimization of lag time underlies antibiotic tolerance in evolved bacterial populations. *Nature* **513**, 418–421 (2014).
2. Venturelli, O. S. *et al.* Deciphering microbial interactions in synthetic human gut microbiome communities. *Mol. Syst. Biol.* **14**, e8157 (2018).
